# DynPerturb:Dynamic Perturbation Modeling for Spatiotemporal Single-Cell Systems

**DOI:** 10.1101/2025.09.15.676236

**Authors:** Hua Qin, Yilin Zhang, Zhihan Guo, Luni Hu, Qingsong Li, Lei Cao, Tianyi Xia, Ziqing Deng, Yong Zhang, Yuxiang Li, Shuangsang Fang

## Abstract

Perturbation responses in multicellular systems are inherently spatiotemporal, governed by temporal dynamics, perturbation intensity, and tissue context. However, existing approaches rarely capture explicit spatiotemporal perturbation dynamics, limiting the ability to reconstruct how perturbations propagate over time, spread across space, and induce feedback at multiple biological scales. Here we introduce DynPerturb, a dynamic inference framework for single-cell and spatial transcriptomics that systematically models the “when-how strong-where” dimensions of perturbations. DynPerturb incorporates temporal encoding, spatial adjacency, and memory modules to capture nonlinear signal propagation and cross-regional feedback with mechanistic interpretability. Across temporal benchmarking datasets, DynPerturb consistently outperformed existing methods in regulatory link prediction, demonstrated zero-shot generalization to unseen nodes, and preserved accuracy under varying sparsity conditions and temporal resolutions. Applied to representative biological systems, DynPerturb uncovers dynamic regulatory rewiring in kidney disease, highlighting a mid-age sensitivity window and potential reversibility with early intervention; delineated a temporal fate-switching boundary between megakaryocyte-erythroid progenitors (MEPs) and granulocyte-monocyte progenitors (GMPs) during hematopoiesis; and revealed spatiotemporal interchamber signaling in murine heart development, where *Igf2* perturbations induced spatial feedback shaping chamber proportioning. Collectively, these results establish DynPerturb as a unified and versatile system for dissecting dynamic perturbation effects and guiding precision intervention strategies in multicellular systems.

## Introduction

Understanding life processes requires modeling regulatory dynamics in real time and space^1^. Without explicit spatiotemporal modeling, analyses often stop at terminal states, making it difficult to identify actionable vulnerable windows and feedback nodes^2–6^. Disease progression, organismal development, and tissue morphogenesis are jointly driven by multilayered regulatory networks unfolding across time and space, where the when (time), how strong (perturbation intensity), and where (space) of perturbation jointly determine outcomes: early interventions can redirect trajectories, whereas late ones face network locking; fine-tuning of perturbation intensity dictates the efficacy-toxicity balance; and the same gene may exert opposite effects in different anatomical regions^2,7–9^. These phenomena essentially reflect the propagation, feedback, and compensation of perturbations in real systems, resulting from the coupling of time-intensity-space (T-I-S). This raises two central challenges for modeling: (i) how to trace perturbation propagation and delayed effects on a real temporal axis, and (ii) how to integrate spatial neighborhoods, tissue mechanics, and intercellular communication into a unified causal framework^1,9,10^.

One established approach reconstructs gene regulatory networks (GRNs) and then performs in silico perturbations on the inferred graph. CellOracle^3^ infers enhancer-resolved regulatory maps from multi-omic data and then applies in silico TF knockout/overexpression to predict state shifts, providing mechanistic interpretability and extrapolation to unseen targets. LINGER^11^ focuses on improving GRN fidelity from paired single-cell RNA-ATAC data and is often paired with CellOracle^3^ for virtual knockouts, thereby increasing confidence in perturbation predictions. GRouNdGAN^12^ embeds user-specified GRNs within a generative framework that simulates realistic single-cell expression and trajectories and supports in silico TF knockouts under non-linear dependencies. Concurrently, temporal models such as dynDeepDRIM^13^ and time-delay LSTM (TDL^14^) improve edge inference on longitudinal single-cell data but rarely encode perturbation intensity or injection timing and seldom enforce causal consistency across time. Despite these advancements, GRN-based simulators are generally limited by predefined or static topologies, which hinder their ability to capture edge rewiring, temporal latency, spatial feedback, and the sensitivity of responses to varying perturbation conditions.

In parallel, another approach involves researchers leveraging single-cell perturbation data to directly learn mappings from cell state and perturbation intensity to perturbation intensity, with increasingly broad capabilities. GEARS^15^ predicts transcriptional responses to unseen single- or multi-gene perturbations at single-cell resolution by learning a deep embedding of perturbation-gene relationships that generalizes combinatorially and across doses. scVIDR^16^ uses a variational autoencoder to model continuous dose-response and interpolate to unobserved doses. CPA^17^ factorizes perturbations into compositional latent variables, unifying dose and combination modeling while retaining interpretability. PRnet^18^ extends this strategy to chemical perturbations by combining molecular representations with conditional generation to predict transcriptional responses of previously unseen compounds. Collectively, this line of work advances perturbation intensity-response modeling, combinatorial perturbations, and cross-batch generalization. However, most models are trained on snapshot data, do not reconstruct time-resolved regulatory processes, and provide limited treatment of causal ordering, latency, and spatial feedback, leaving a clear opportunity for unified modeling across time, intensity, and space.

To bridge this gap, we present DynPerturb, a unified dynamic pertubation modeling framework for real-time, perturbation intensity, and spatial (T-I-S) coupling. Distinct from “static network and endpoint prediction” approaches, DynPerturb directly learns the propagation, reconstruction, and feedback of perturbations on real-time graphs. It integrates time, perturbation intensity, and spatial information into a unified representation, enabling structured inference of regulatory responses to localized and temporally varying perturbations across biological systems. This is achieved by modeling the timing of external signals, their varying intensity, and site-specific interactions within dynamic gene regulatory networks. This design allows simultaneous tracking of causal propagation, intensity-dependent modulation, and spatially resolved feedback within a single computational framework.

We validate the methodology of DynPerturb by benchmarking across multiple real time-series datasets, evaluating DynPerturb’s performance in dynamic GRN reconstruction, zero-shot extrapolation, and hyperparameter sensitivity, and confirming robustness to sample size, feature selection, and temporal resolution through ablation analyses. We then demonstrate three end-to-end applications of DynPerturb: (i) reconstructing time-dependent networks in chronic kidney disease to identify high-responsiveness windows and early damage-repair trajectories; (ii) recapitulating directional lineage bias driven by seven hematopoietic transcription factors, and dissecting combinatorial and time-dependent fate shifts; and (iii) integrating spatial transcriptomics in mouse heart development to reveal region-specific signaling dynamics and tissue patterning during ventricle-atrium formation. Collectively, DynPerturb transforms “when to intervene, how strong to apply, and where to target” into testable, quantitative, and interpretable hypotheses and strategies, offering a unified modeling pathway from perturbation input to system response under the T-I-S coupling perspective.

## Results

### Overview of the DynPerturb Framework

DynPerturb serves as a unified dynamic perturbation modeling framework, seamlessly integrating multimodal data fusion, graph construction, spatiotemporal encoding, and perturbation injection into a cohesive workflow (**Fig. 1A**). Specifically, it incorporates gene expression matrices alongside auxiliary biological features, such as regulatory dependencies, developmental trajectories, and spatial adjacency data, to capture comprehensive system-level information. At each discrete time point, a dynamic graph is constructed to delineate the underlying biological structure: nodes in this graph represent either cells or genes, while edges encode regulatory interactions (e.g., gene-gene regulation) or spatial associations (e.g., cell-cell proximity). Notably, all temporal graphs share a unified node set and are embedded within a common latent space; this design enables the model to quantitatively monitor both structural remodeling (e.g., edge rewiring) and state transitions (e.g., cell phenotype shifts) induced by external perturbations.

**Figure 1.**
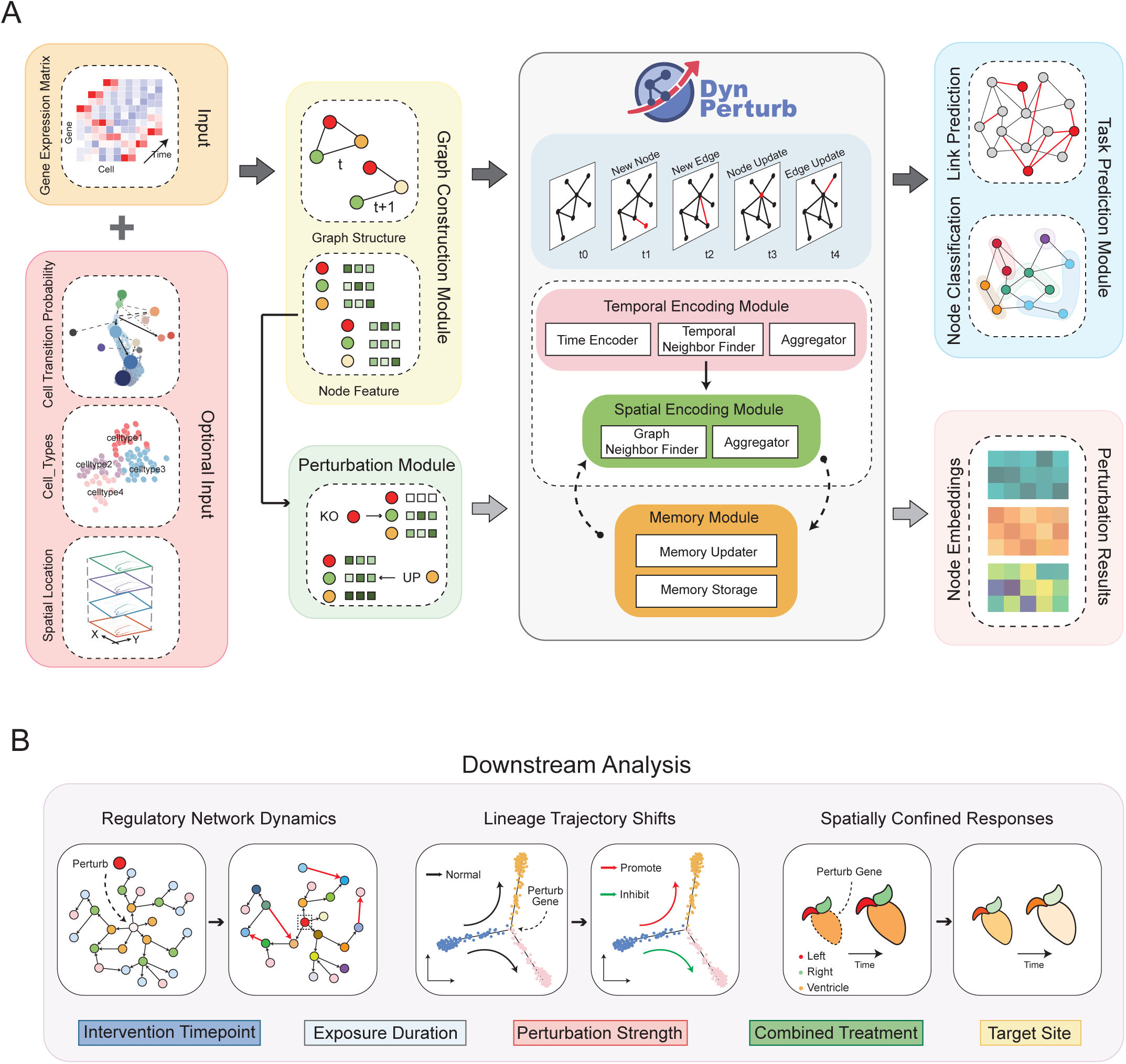
DynPerturb enables dynamic perturbation modeling. **(A)** Overview of the DynPerturb framework. Single-cell data across time points are integrated into dynamic graphs with temporal encoding and memory modules, supporting downstream prediction tasks and perturbation simulation. **(B)** Example applications of DynPerturb showing predicted effects of gene perturbations at the gene regulatory, cell fate, and spatial patterning levels.The model can incorporate various perturbation factors, including intervention timepoint, exposure duration, perturbation strength, combined treatment, and target site.

To accurately capture the spatiotemporal trajectory of perturbation propagation, DynPerturb integrates joint temporal and spatial encoding modules. The temporal encoding module is built on time-aware graph neural networks, which fuse three core components: time encoders, adaptive neighborhood selection mechanisms, and temporal aggregation strategies, to learn the history-dependent propagation patterns of perturbations. This allows the model to explicitly model how perturbation effects evolve dynamically over time, accounting for prior system states. Concurrently, the spatial encoding module leverages anatomical context within graph neighborhoods to construct fine-grained local response representations. These representations effectively capture region-specific feedback effects (e.g., tissue microenvironment-dependent responses) and intra-tissue heterogeneity, providing critical spatial dimensionality for characterizing node dynamics.

The fused spatiotemporal representations collectively determine the dynamic state of each node, which is then transmitted to a dedicated memory module. This module maintains time-stamped hidden states for individual nodes; drawing inspiration from **Temporal Graph Networks** (TGN^19^), it facilitates the modeling of long-range temporal dependencies and nonlinear feedback loops while enforcing temporal consistency. This design ensures reliable reconstruction of perturbation trajectories within the target biological system. To comprehensively characterize perturbed system dynamics, DynPerturb jointly optimizes two complementary tasks: link prediction (reconstructing latent regulatory network rewiring, e.g., gene regulatory edge gain/loss) and node classification (tracking cell state evolution, e.g., transcriptional reprogramming). It accommodates multi-source supervision (known cell annotations, lineage trajectories, spatial labels), enhancing interpretability of learned biological mechanisms and generalization to unseen perturbations.

This framework is specifically tailored for modeling complex biological processes, such as embryonic development, tissue injury response, and regenerative repair. In DynPerturb, perturbations are represented as flexible, time-stamped input events, with configurable parameters that control the intervention timepoint, exposure duration, perturbation intensity, combined treatment, and target site, enabling detailed modeling of regulatory network dynamics, lineage trajectory shifts, and spatially confined responses (**Fig. 1B**). These perturbation inputs can be injected into arbitrary nodes within the graph, triggering cascading local responses and system-wide effects via dynamic graph updates. Collectively, these components enable DynPerturb to simulate the propagation, compensation, and feedback of perturbations within a causally consistent spatiotemporal framework, providing a generalizable platform for modeling key biological phenomena such as transcriptional reprogramming, tissue-level heterogeneity, and regulatory network destabilization across diverse biological systems.

### Advantages and generalization capacity of DynPerturb in modeling dynamic regulatory networks

To assess the capacity of DynPerturb for modeling dynamic regulatory interactions, we performed link prediction across four public time-series datasets^14^, comprising two human embryonic stem cells (hESCs) and two mouse embryonic stem cells (mESCs) systems (**Supplementary Fig. 1D**). DynPerturb achieved an overall average AUC of 0.892 ± 0.024 (n=4), consistently outperforming representative baseline methods across modeling paradigms (**Fig. 2A**), with relative improvements of approximately 22%-51% (vs. TDL/dynDeepDRIM ≈ 22%, vs. PCC/MI ≈ 44%-51%) (**Supplementary Table 1**). We excluded CellOracle^3^ from the baseline comparisons because it relies on static gene regulatory networks, which are unable to capture temporal dynamics. Similarly, GEARS^15^ was not included, as it is trained on snapshot perturbation data and does not reconstruct time-resolved causal regulatory processes, making it fundamentally different from the temporal modeling task addressed by DynPerturb. Ablation analysis (**Fig. 2B;Supplementary Fig. 1B**) demonstrated that DynPerturb is robust to variations in sample size and the dimensionality of node features (defined by the number of highly variable genes), and exhibits expected monotonic gains with increased temporal resolution. Specifically, increasing the number of observation time points from 3 to 9 yielded AUC gains from 0.808 to 0.916/0.915 (Δ ≈ +0.108), plateauing around ≥ 7 time points. These results support the generalizability and robustness of DynPerturb across diverse levels of network sparsity and dynamical regimes. A core advantage of DynPerturb lies in its zero-shot link prediction for unseen nodes(**Supplementary Fig. 1C**), which is facilitated by temporally conditioned encodings and inductive message passing mechanisms that extrapolate from node features and real-time context. Across all four datasets, DynPerturb outperformed random baselines by an average of +22.7 percentage points in zero-shot AUC (**Fig. 2C**).

**Figure 2.**
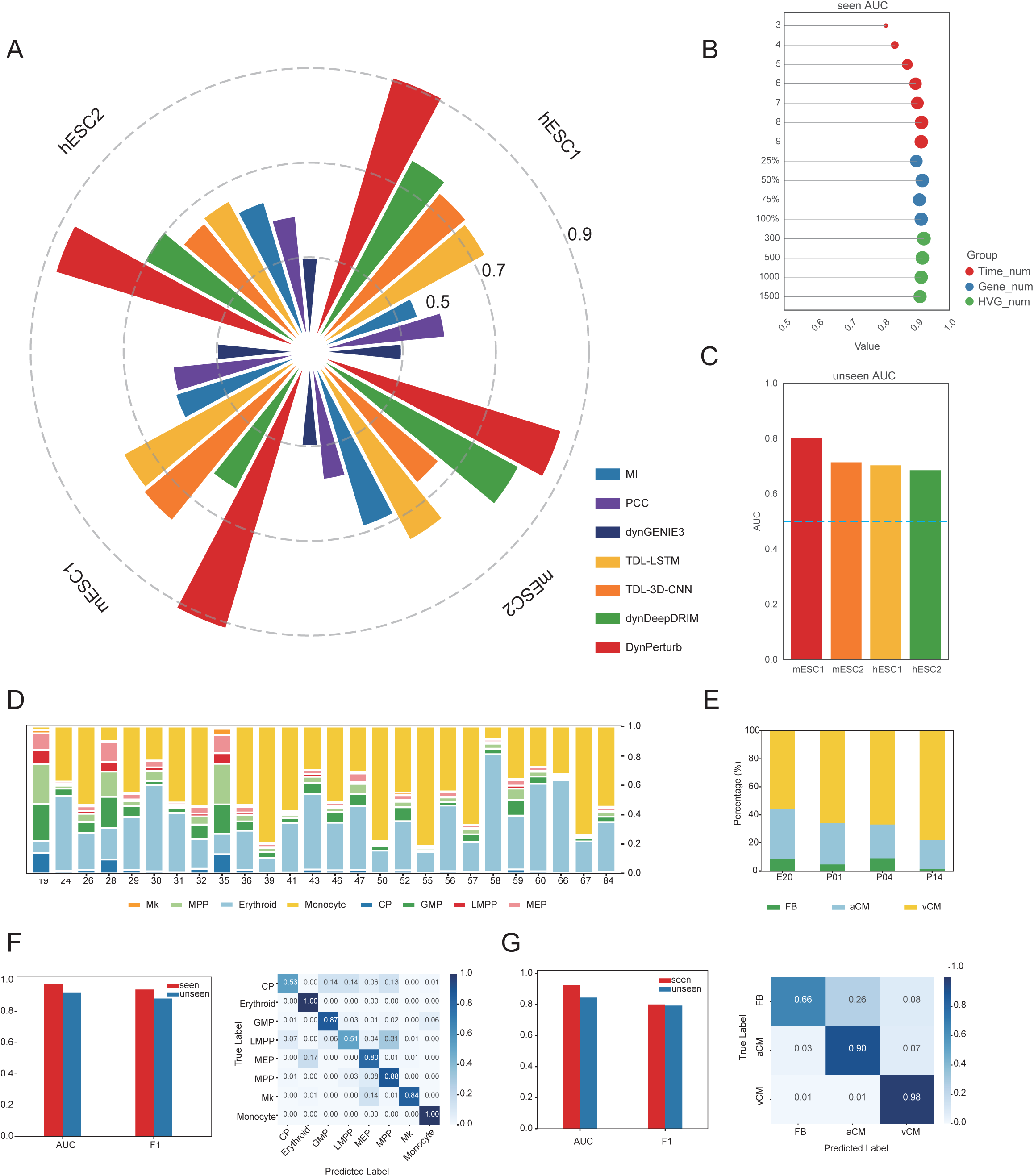
Performance evaluation of DynPerturb on link prediction and node classification tasks. **(A)** Comparison of link prediction AUC between DynPerturb and six baseline methods across four benchmark datasets. **(B)** Ablation study on the number of time points, gene numbers, and highly variable gene (HVG) numbers for link prediction AUC on seen nodes in the mESC1 dataset. **(C)** Comparison of link prediction AUC on unseen nodes across four benchmark datasets. **(D)** Cell type composition across time points in the hematopoietic dataset. **(E)** Proportions of different cell types across the four developmental stages. **(F)** Evaluation of link prediction (AUC) and node classification (confusion matrix) of Hematopoiesis dataset. **(G)** Evaluation of link prediction (AUC) and node classification (confusion matrix) of Cardiomyocyte differentiation dataset.

We also introduced a biologically supervised branch for node classification, such as cell identity or anatomical annotation, which is jointly trained with link prediction, to align regulatory topology with phenotypic boundaries. Applied to a 26-timepoint human hematopoiesis dataset spanning ages 19 – 84 (**Fig. 2D;Supplementary Fig. 7**), and to a spatial transcriptomic atlas of the mouse heart across postnatal stages (E20/P01/P04/P14; vCM/aCM/FB) (**Fig. 2E;Supplementary Fig. 17B-C,E**), multi-task DynPerturb achieved AUCs > 0.9 in link prediction, while the node classification confusion matrices showed strong diagonal patterns. In hematopoiesis, the classification performance is highly accurate with an average accuracy of **0.958**, while in the heart atlas, the average accuracy is **0.956** (**Fig. 2F-G**). Thus, the cell-annotation task sharpens phenotypic boundaries while preserving link-prediction performance.

### Temporal perturbation mapping reveals regulatory fragility in CKD tubule cells

To evaluate whether DynPerturb can faithfully capture the dynamic remodeling of gene regulatory networks along the trajectory of disease progression and simulate perturbation propagation across heterogeneous cell populations under different perturbation scenarios, we applied DynPerturb to a single-cell transcriptomic dataset encompassing 7,853 human proximal tubule (PT) cells and abnormal proximal tubule (aPT) cells (**Fig. 3A**). These samples span 15 chronological stages (ages 30-66) (**Supplementary Fig. 3A-C**) and were derived from patients with chronic kidney disease (CKD) and acute kidney injury (AKI).Cells were annotated into nine distinct subpopulations, ranging from early progenitor-like subsets (aPT-A/B) cells with regenerative potential to late-stage degenerative states (PT-S2a/b) (**Fig. 3B**). DynPerturb reconstructed dynamic gene regulatory networks (GRNs) for each subpopulation at each time point, allowing longitudinal tracking of transcriptional states and identification of perturbation-specific response trajectories.To systematically interrogate key regulators underlying the transition from AKI to CKD, we focused on three key transcription factors, *ELF3*, *KLF6*, and *KLF10*, that were consistently upregulated in aPT cells (**Fig. 3C; Supplementary Fig. 3E**,5**-6**).

**Figure 3.**
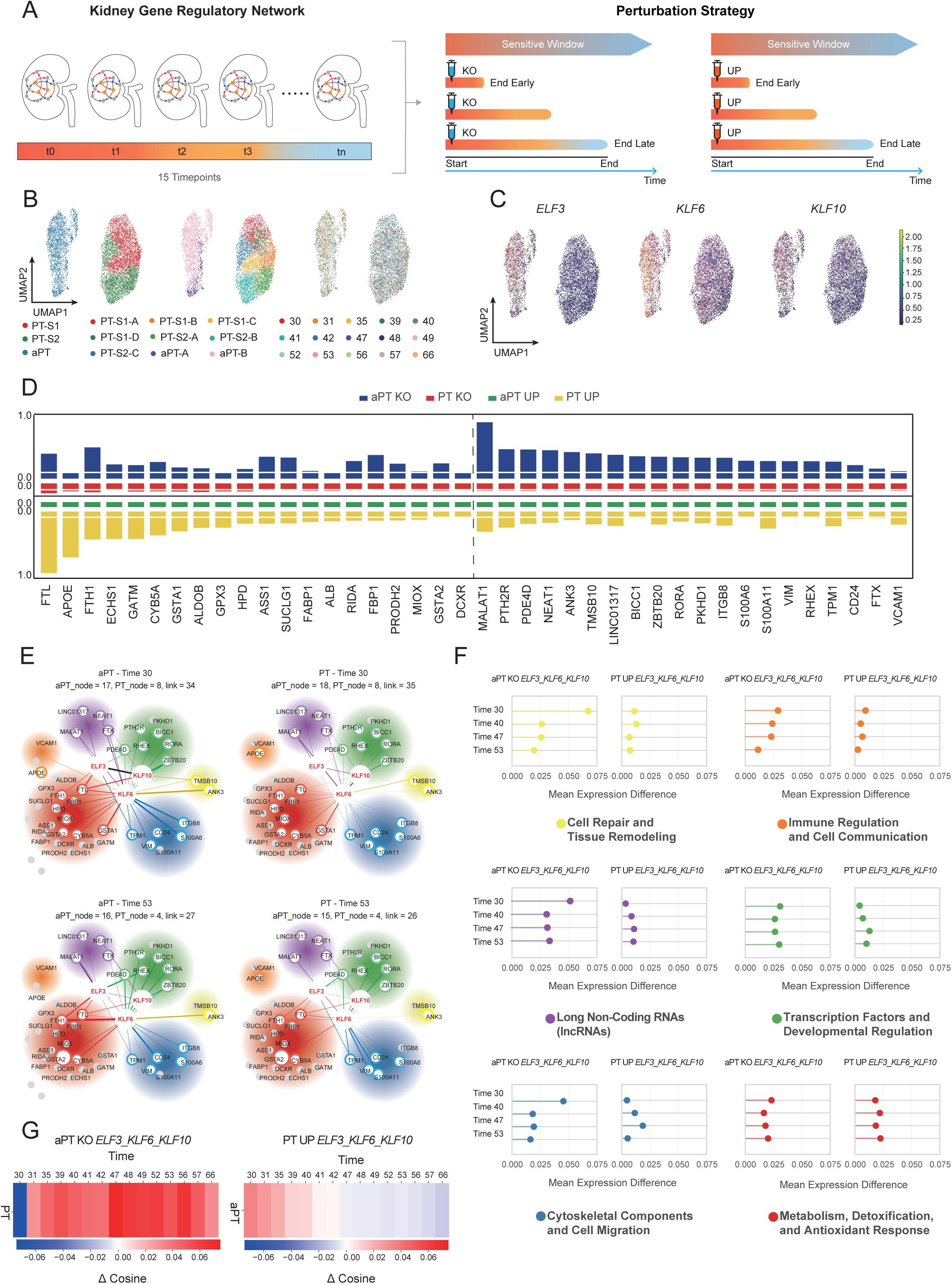
TF KO/UP simulations reveal tubular fate shifts and sensitive window. **(A)** Dynamic modeling of proximal tubule regulatory networks and perturbation strategies through simulated knockout (KO) and overexpression (UP) of key transcription factors. **(B)** UMAP of cells from diseased kidneys, colored by celltype, cluster, and time point. **(C)** Expression patterns of *ELF3*, *KLF6*, and *KLF10* across the dataset. **(D)** Absolute changes in node embeddings of 40 marker genes (20 PT markers and 20 aPT markers, separated by a dashed line) after perturbing *ELF3*, *KLF6*, and *KLF10* in both PT and aPT. Each row corresponds to one perturbation condition (aPT KO, PT KO, aPT UP, or PT UP) across multiple time points. **(E)** Gene regulatory network (GRN) diagrams of *ELF3*, *KLF6*, *KLF10*, and the 40 markers at time points 30, 40, 47, and 53. *ELF3*, *KLF6*, and *KLF10* are shown in red; PT markers are in blue shades, aPT markers are in green shades; deeper colors indicate larger changes in node embeddings; edge thickness represents regulatory strength; gray nodes indicate genes absent in the GRN at that time point. **(F)** Lollipop plots summarizing total node embedding changes of the 40 markers grouped into six functional categories at time points 30, 40, 47, and 53. Left: aPT KO condition; right: PT UP condition. **(G)** Changes in cosine distance between PT and aPT over time after perturbing *ELF3*, *KLF6*, and *KLF10*. Left: aPT KO (aPT to PT direction); right: PT UP (PT to aPT direction). Red indicates a healthier state (more PT-like), whereas blue indicates a more diseased state (more aPT-like).

#### DynPerturb-mediated phenotypic reprogramming and dynamic regulatory remodeling of proximal tubule cells mediated by *ELF3/KLF6/KLF10*

We assessed four perturbation scenarios involving either sustained overexpression or knockout of *ELF3*, *KLF6*, and *KLF10* in both PT and aPT cells (**Fig. 3D; Supplementary Fig. 4A**). These scenarios included aPT-KO (knockout in diseased state), aPT-UP (overexpression in diseased state), PT-UP (overexpression in healthy state to mimic pathological induction), and PT-KO (knockout in healthy state to test homeostatic dependency). Under the aPT-KO condition, DynPerturb predicted marked shifts in the expression of aPT-associated genes (e.g., *MALAT1*, *NEAT1*, *TMSB10*), along with restoration of PT marker gene expression (e.g., *GSTA1*, *HPD*, *FABP1*) (**Fig. 3D**, blue bars; **Supplementary Fig. 3D**). These coordinated changes suggest a transcriptional “depathologization” of aPT cells. Supported by previous spatial multi-omic and animal model evidence^20–22^, non-cell-autonomous mechanisms such as paracrine signaling, cell – cell contact, and matrix remodeling may amplify repair signals and influence neighboring cells. In contrast, sustained overexpression of *ELF3/KLF6/KLF10* in PT cells (PT-UP) disrupted the homeostatic baseline, resulting in fluctuations in PT markers (e.g., *FABP1*, *ECHS1*, *GSTA1*) and activation of pathological effectors such as *VCAM1*^23,24^ and *ITGB8*, shifting the transcriptomic state toward an aPT-like phenotype (**Fig. 3D**, yellow bars). Notably, neither aPT-UP nor PT-KO induced substantial phenotypic changes, underscoring the state-dependent and the inertia-prone behavior of the regulatory network (**Supplementary Fig. 4B**).

To explore the downstream fate consequences of these perturbations, we reconstructed *ELF3/KLF6/KLF10*-centered gene regulatory networks in 30- and 53-year-old samples via DynPerturb (**Fig. 3E**). Both aPT-KO and PT-UP exhibited network simplification, reflected by reductions in node and edge counts. The effect was more pronounced in PT-UP (−26.9% nodes, −25.7% edges) than in aPT-KO (−20.0% nodes, −20.6% edges), indicating a more pronounced topological collapse when homeostasis is disrupted. To interpret the biological implications of network contraction, we functionally categorized 40 core target genes into six modules: metabolism/antioxidation, cytoskeleton, developmental regulation, long non-coding RNAs(lncRNAs), immune modulation, and tissue remodeling, and meanwhile quantified perturbation magnitude across time (**Fig. 3F**). Under aPT-KO, a high-initial-perturbation followed by gradual resolution dynamic emerged. Our results showed that structural modules such as cytoskeleton, lncRNAs, and remodeling exhibited strong early perturbation followed by convergence, consistent with prior reports^25^. In contrast, emetabolic and immune modules displayed delayed recovery, also in line with previous findings. Developmental regulators remained relatively stable at intermediate stages, supporting earlier observations^26^ that prolonged signal integration is required during fate reprogramming. By contrast, PT-UP triggered weaker yet chronically accumulating perturbations across modules, following a “slow-drift” trajectory that gradually diverted cells from their steady-state toward subclinical pathological transitions. This pattern recapitulates canonical disease kinetics, in which early compensation masks symptoms until a critical threshold is breached^27^. In such contexts, targeted therapies may restore system balance more rapidly by interrupting causal axes earlier in the process.

#### DynPerturb reveals transcription factor imbalance drives proximal tubule cell fate shifts and identifies a mid-life vulnerability window

Based on the aforementioned perturbation trends, we further employed DynPerturb to trace and assess systemic fate transitions under distinct intervention conditions (aPT-KO and PT-UP) (**Fig. 3G; Supplementary Fig. 4C**), using graph-level embeddings derived from GCN (**Methods**). It was discovered that under the aPT-KO condition, the deletion of these transcription factors (TFs) prompted aPT cells to transition towards aPT-like phenotype as early as 30 years of age. This transition was accompanied by a diminishment of the characteristics associated with inflammatory factors (*VCAM1*), structural remodeling factors (*TMSB10*), and fibrotic lncRNAs (*MALAT1*, *NEAT1*), suggesting that cells maintained plasticity and phenotypic reversibility once the pathological state was alleviated. This finding has been corroborated in relevant studies^28–30^. Conversely, PT-UP did not instigate acute perturbations in the early stages. Instead, it gradually shifted the trajectories towards the pathological state trajectory commencing at 40 years of age, with a progressive accumulation of perturbations in immune and structural modules. This indicates that chronic stimulation was disrupting the original homeostasis^23^. Particularly during the mid-life sensitive windows (42-56 years), both cell types exhibited synchronous and significant alterations in their trajectories, indicating that this stage may represent a potential window for intervention response.

#### Directional and temporal in-silico knock-out experiments reveal regulatory inflection time points in CKD progression

To assess the ability of DynPerturb to predict the impact of perturbations applied at different time points, we simulated the dynamic responses of aPT-A/B subpopulations to either knockout or upregulation of *ELF3*, *KLF6*, and *KLF10* at ages 30, 39, 42, 49, and 56 (**Fig. 4B; Supplementary Fig. 6B**), aiming to identify potential temporal windows for aPT cell-state recovery and age-dependent regulatory inflection points. The predictions revealed that aPT-B, compared with aPT-A, exhibited clearer response directionality and greater sensitivity, indicating higher transcriptional plasticity^23^, thus, subsequent analyses focused on aPT-B. Under knockout conditions, transcriptional reprogramming was markedly induced at ages 30 and 39, reflecting preserved system responsiveness. In contrast, responses were substantially attenuated at ages 49-56, suggesting network desensitization-a finding consistent with independent evidence for age-related decline in renal repair capacity^31^. In comparison, sustained TF upregulation at all ages failed to elicit notable gene expression changes, indicating that in the CKD background, indicating the system may already be in a state of activation saturation and is more sensitive to deactivation^23^, reflecting a high-tension, edge-unstable state.

**Figure 4.**
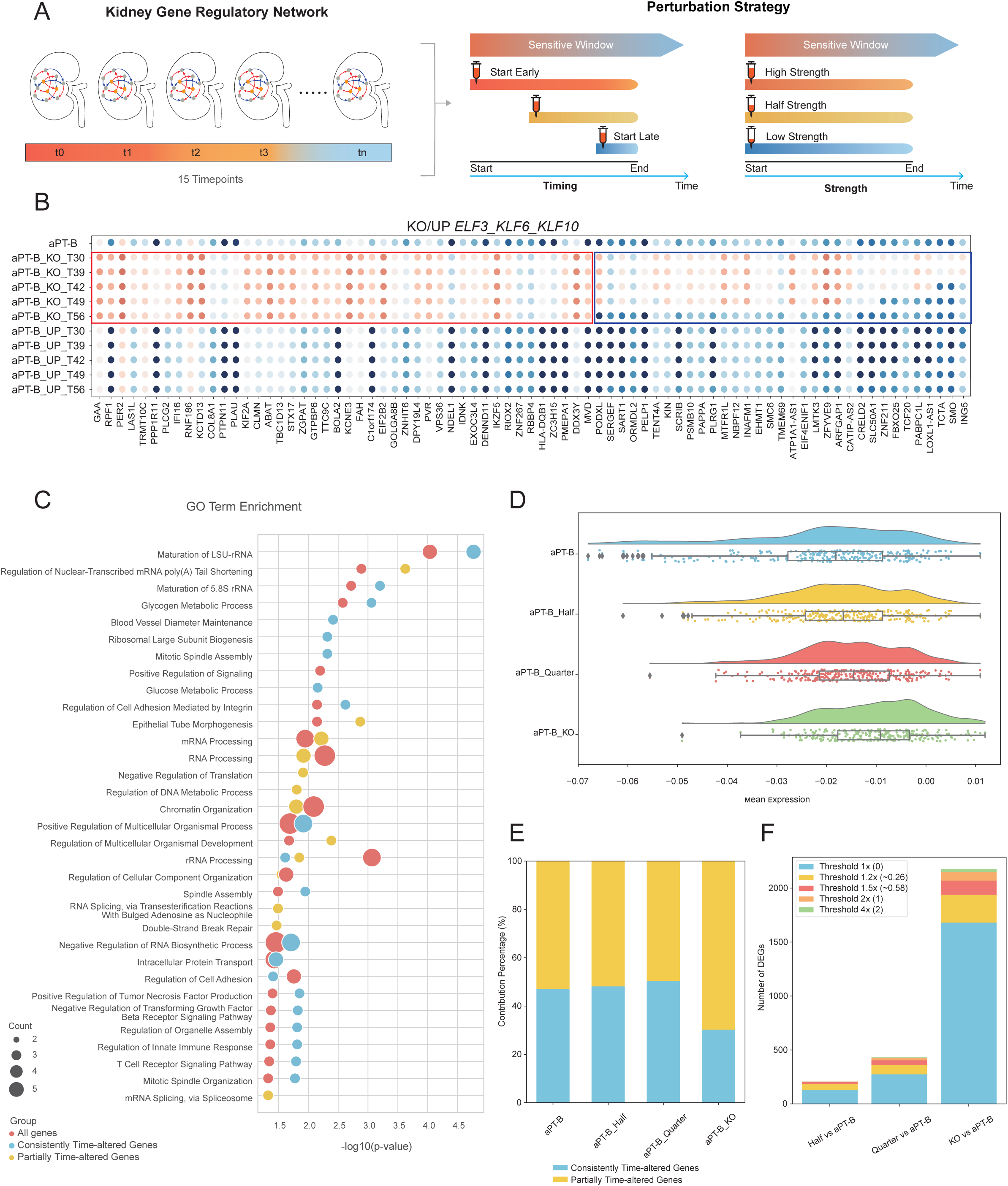
Timing and perturbation strength simulations uncover temporal sensitivity and nonlinear tipping points in CKD regulatory networks. **(A)** Dynamic modeling of proximal tubule regulatory networks and perturbation strategies across time and intensity. **(B)** Embedding dynamics following knockout (KO) or upregulation (UP) of *ELF3*, *KLF6*, and *KLF10* starting at different time points in aPT-B. The 80 most perturbed genes are shown; red-boxed genes exhibit similar changes regardless of the knockout timing, and blue-boxed genes show early knockout-specific changes with minimal late knockout effects. The effect of upregulation on aPT-B node embeddings is negligible. **(C)** Gene Ontology (GO) enrichment analysis for the three groups: All genes, Consistently Time-altered Genes, and Partially Time-altered Genes. **(D)** Node embedding changes after full, quarter (reduced to one-quarter of the original expression level), or half knockout of *ELF3*, *KLF6*, and *KLF10* from the earliest time point, visualized using the top 80 perturbed genes. Quarter represents perturbation to one-quarter of the original expression level. The distribution is shown as a raincloud plot. **(E)** Contribution proportion of Consistently Time-altered Genes and Partially Time-altered Genes under the four KO conditions shown in (C). **(F)** Stacked bar plots showing the number of differentially expressed genes (DEGs) compared to the original aPT-B under different thresholds across three knockout strengths (full, quarter, and half).

We further categorized responsive genes into “persistent responders” and “early responders with late silencing”, and integrated these profiles with system-level functional enrichment analysis. The results indicated a temporally ordered activation across three functional axes^23,31^ (**Fig. 4C**). Genes with early but transient responses were mainly enriched in the immune surveillance axis, such as *IFI16* and *HLA-DQB1*, which were rapidly downregulated within the *NF-κB* and *C-type lectin receptor* pathways, suggesting early decoupling of inflammatory signaling^32^. The structural homeostasis axis showed transient regulatory adjustments, including tension relief and secretion regulation genes such as *COL8A1* and *GOLGA8B*. In contrast, the fate-regulatory axis (e.g., *ZNF267*, *SMO*) displayed only mild and transient compensatory activation that quickly subsided over time. These findings indicate that TF knockouts trigger a transient system-level response in which pathway accessibility initially decreases and ultimately collapses. The resulting trajectory reveals a reversible intervention window during which regulatory modules can be temporarily uncoupled, structural adjustments occur, and lineage commitments remain unstable, consistent with prior reports that, without timely intervention, the system drifts from plasticity to an irreversibly restructured steady state^31^.

#### Perturbation strength-dependent modeling reveals nonlinear thresholds and network stability in aPT-B

For the purpose of evaluating the sensitivity of DynPerturb to perturbation strength, we simulated transcription factor knock-out gradients (50%, 75%, 100%) of *ELF3*, *KLF6*, and *KLF10* at the 30-year-old timepoint using the DynPerturb predictive framework. This enabled us to observe how the key CKD subpopulation aPT-B responds to varying intensities of perturbation, and to trace the trajectories of the 80 most responsive genes within the learned embedding space (**Fig. 4D; Supplementary Fig. 4D**). Displacement within the embedding indicates global rewiring of the regulatory network^3,33^, consistent with previous observations. However, this trend underwent a nonlinear transition, where the system stayed stable through redundancy and localized compensatory mechanisms under low to moderate perturbation strengths, but multiple functional modules collapsed simultaneously once the perturbation strength went beyond the critical threshold (∼75%). Under full-strength combinatorial knock-out, maximal gene displacement was observed, indicating that the system had crossed a homeostatic limit into global reprogramming^34^. Quantitative analyses (**Fig. 4E-F**) showed a sharp increase in both the number of responsive genes and the magnitude of their displacement beyond the 75% threshold, indicating a phase transition-like destabilization. This nonlinear, threshold-dependent response is quantitatively reproduced by DynPerturb, enabling prediction of latent collapse windows for perturbation-strength-specific interventions.

### Lineage-specific transcription factor perturbations shape hematopoietic trajectories

To evaluate DynPerturb’s capacity to detect and quantify transcriptional perturbations across lineage development (**Fig. 5A**), we applied the model to a human bone-marrow dataset comprising 26 time points from individuals aged 19 to 84 years (**Fig. 5B**). We then constucted the developmental trajectory (**Fig. 5C**) to chart the hematopoietic differentiation timeline and facilitate in silico perturbation assays. Seven transcription factors with well-established lineage directing roles, including *IRF8*, *CEBPA*, *CEBPE*, *GATA1*, *KLF1*, *GATA2*, and *FLI1*, were selected for virtual knock-out experiments to systematically evaluate their effects on megakaryocyte-erythroid (ME) and granulocyte-monocyte (GM) differentiation trajectories (**Fig. 5D-E**). By leveraging directionality metrics derived from vector fields (Perturbation Score) (**Methods**) alongside quantitative estimates of perturbation magnitude, DynPerturb effectively captured lineage redirection tendencies, stage-dependent sensitivities, and perturbation strength-dependent responses induced by transcriptional perturbations (**Fig. 5F**). Many of the predicted responses were validated and were consistent with prior biological knowledge. Specifically, knock-out of *GATA1* or *KLF1* markedly impaired ME differentiation, while deletion of *IRF8* or *CEBPA* substantially disrupted GM development. The effects of other factors aligned with established regulatory functions^3,35,36^, consistent with previous reports. Complementary simulations of sustained overexpression promoted lineage-biased differentiation consistent with the known roles of these transcription factors (**Supplementary Fig. 8-11**).

**Figure 5.**
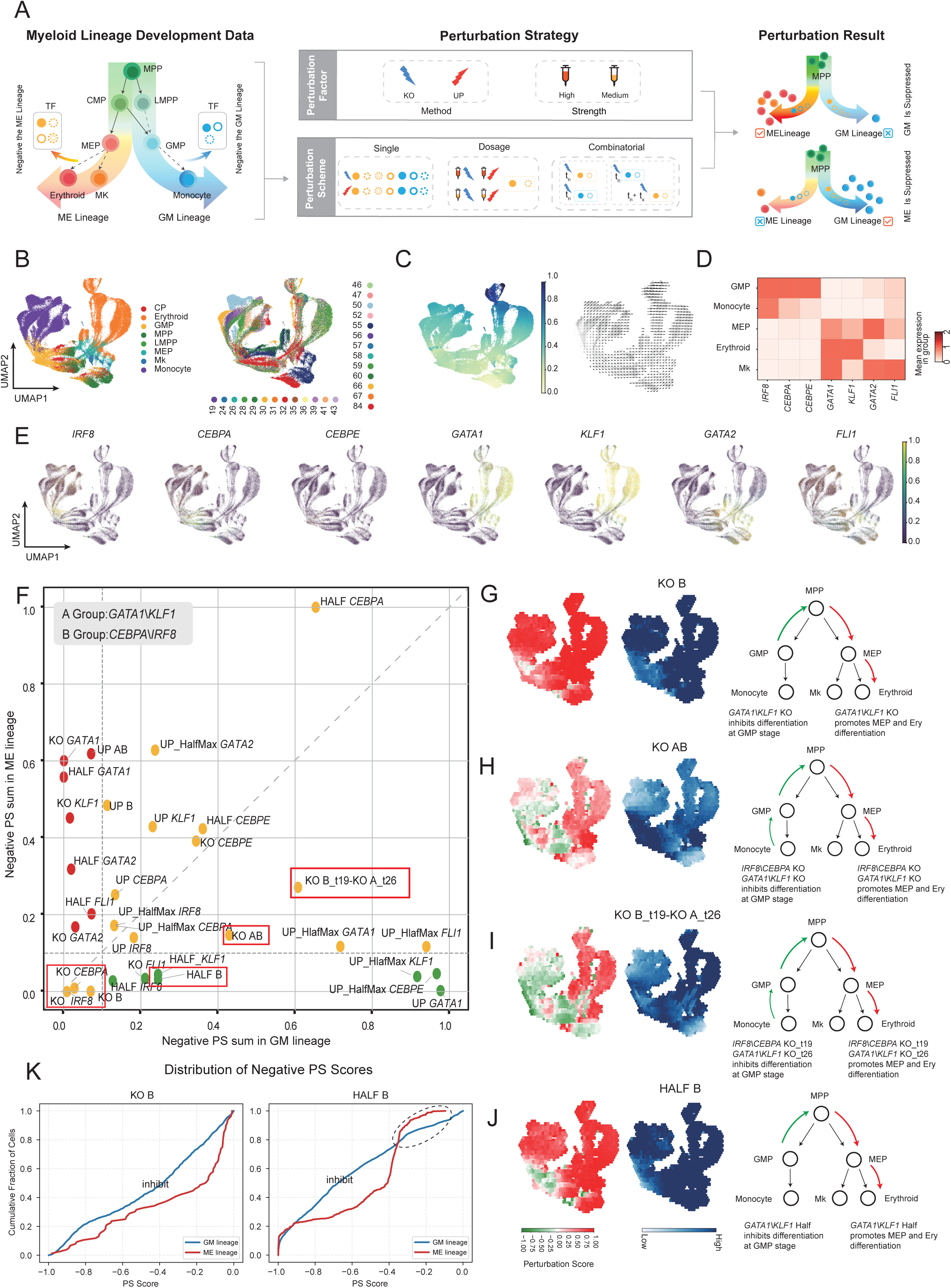
Cell-level perturbations disclose lineage-specific regulatory sensitivity during human hematopoiesis. **(A)** Perturbation analysis of myeloid lineage development, showing developmental data, perturbation factors and strategies including method, strength, and combinations, and the resulting effects on lineage progression. **(B)** UMAP of human bone marrow cells colored by cell type and time point. **(C)** Pseudotime trajectory inferred by Palantir and its corresponding developmental vector field. **(D)** Expression highlights of the seven regulators across the dataset. **(E)** Differential effects of seven key transcription factors along ME and GM lineages. **(F)** Scatterplot showing negative perturbation scores (PS) for all perturbations across erythroid and GM lineages. Proximity to axis indicates stronger suppression of that lineage. **(G)** Effects of knocking out B-group genes (*CEBPA*, *IRF8*). **(H)** Effects of simultaneous knockout of both A-group (*GATA1*, *KLF1*) and B-group (*CEBPA*, *IRF8*) genes. **(I)** Sequential knockout: B-group genes at time 19 and A-group genes at time 26. **(J)** Strength effect: Half knockout of B-group genes. **(K)** Cumulative distributions of negative perturbation scores (PS) for GM and ME lineages under full (KO_B) and partial (HALF_B) knockout conditions. In both settings, GM exhibits a broader distribution of negative PS values, indicating stronger suppression. Under partial knockout (right), this suppression is mitigated, and the ME lineage shows a relative increase in low PS values, reflecting differential sensitivity to perturbation intensity.

#### DynPerturb reveals severe suppression of GM lineage upon simultaneous knockout of GM factors

We next examined the functional interplay between two key transcription factors, *IRF8* and *CEBPA*, within the granulocyte-monocyte (GM) lineage(**Supplementary Fig. 14A**). *IRF8* primarily acts at the granulocyte-monocyte progenitor (GMP) stage to promote the transcriptional activation of monocyte and dendritic-like fates, while *CEBPA* facilitates the differentiation of multipotent progenitors (e.g., CMPs) toward GMP and initiates granulocytic programs^37,38^. DynPerturb simulations revealed that concurrent deletion of *IRF8* and *CEBPA* in 19-year-old samples led to a collapse of GM lineage development: progenitor cells failed to progress along either the GMP or monocyte path and instead exhibited a pronounced shift toward the megakaryocyte-erythroid (ME) lineage (**Fig. 5F-G**). This effect could not be explained by additive contributions alone, but rather reflects a regulatory interdependence between the two factors, highlighting a non-redundant bottleneck in GM specification and an asymmetric plasticity of lineage fate^39,40^.

Mechanistically, *CEBPA* normally maintains GM identity by repressing erythroid regulators such as *GATA1* and *KLF1*, while *IRF8* prevents nonspecific deviation of granulocytic trajectories toward erythroid fates. Upon simultaneous deletion, these differentiation constraints are lost, allowing the erythroid program to be derepressed and driving progenitor cells to default toward the ME lineage^39,40^. Through this process, DynPerturb not only reconstructed the asymmetric plasticity inherent in key regulatory axes, but also elucidated the underlying structural dependencies of the transcriptional network that govern lineage redirection.

#### Fate Restoration of GM Lineage and Plasticity Window under Competitive Lineage-Coupled Regulation

To dissect the competitive dynamics underlying lineage fate regulation and examine how these systems respond to perturbations, we utilized DynPerturb for a dual knock-out simulation targeting GM-promoting (*IRF8*, *CEBPA*) and ME-promoting (*GATA1*, *KLF1*) transcription factors(**Supplementary Fig. 14B**). When compared with GM-specific deletion that caused a unidirectional drift toward ME identity (**Fig. 5G**), concurrent disruption of both lineage regulators resulted in partial lineage restoration, most notably the re-establishment of the GMP branch (**Fig. 5F,5H**). The perturbation score within the GM compartment shifted from strongly negative toward neutrality, indicating the release of latent GM differentiation capacity once ME inhibition was lifted. These results highlight a mutual antagonism between the GM and ME programs. Supporting this mechanism, prior studies have demonstrated that *GATA1/KLF1* downregulate *PU.1* and *IRF8* to block GM specification, whereas *IRF8* reciprocally restricts *GATA1*-driven erythroid differentiation^41,42^. Removal of both sets of regulators eliminated this mutual inhibition, creating a regulatory void in which hybrid lineage potentials emerged and compensatory shifts toward GMP commitment were initiated.

We next explored the influence of intervention timing on GM lineage recovery by implementing a temporally staggered perturbation: GM factor deletion (*IRF8*, *CEBPA*) was introduced at age 19, followed by ME factor deletion (*GATA1*, *KLF1*) at age 26 (**Fig. 5F,5I**). According to previous studies^41,42^, despite early suppression of GM programming, a subset of progenitor cells retained the capacity for subsequent trajectory re-engagement, resulting in partial GMP recovery. This behavior suggests the existence of a “plasticity window” during which lineage commitment remains conditionally reversible, reinforcing the importance of temporal context in shaping fate decisions.

DynPerturb successfully reconstructed the elastic remodeling of hematopoietic fate trajectories under these layered perturbations, indicating the central role of non-linear regulatory coupling in maintaining system resilience and coordinating responses to disruption^43^. Quantitative mapping of perturbation timing and signal integration by the DynPerturb framework establishes lineage fate as a time-dependent readout, yielding precise intervention windows aligned with prior reports^35,37^.

#### Gradual but stable lineage bias induced by perturbation-strength-dependent modulation

To assess DynPerturb’s capacity to capture responses under varying perturbation strength was conducted by simulating a 50% knockdown of the GM-regulating transcription factors *IRF8* and *CEBPA* (**Fig. 5F,5J;Supplementary Fig. 12-13,15-16**). Compared to complete knockout (**Fig. 5G**), suppression of the GM branch was substantially mitigated, with perturbation scores distributed more evenly, indicating a milder and more stable lineage shift. The ME branch exhibited slight enhancement, though less pronounced than under full knockout, suggesting nonlinear responsiveness of the regulatory network to perturbation strength.This nonlinear response was further evidenced by changes in the empirical cumulative distribution of negative perturbation scores (**Fig. 5K**). Under complete knockout (KO_B), the empirical cumulative distribution function (ECDF) curve for the GM lineage shifted markedly leftward, indicating stronger suppression(**Methods**). In contrast, under partial knockout (HALF_B), the ECDF curves for GM and ME lineages became more aligned, and the tail of the ME curve rose, reflecting partial recovery of lineage-associated expression. Unlike the lineage collapse induced by full perturbation, the GM branch retained regulatory pathway integrity under partial knockdown, suggesting the system remained in a metastable, transcriptionally plastic state.These findings demonstrate that GM differentiation progresses through a perturbation strength governed continuum. Partial retention of *IRF8* and *CEBPA* activity supports maintenance of the core developmental trajectory. This sub-threshold steady state reflects the robustness of developmental systems, whereby moderate perturbations are buffered to preserve lineage fidelity and prevent erroneous fate transitions, consistent with previous studies ^39,40^. DynPerturb effectively delineates how changes in perturbation magnitude influence the propagation of regulatory effects and lineage deflection, positioning perturbation sensitivity as a critical variable in hematopoietic decision-making and offering a rational foundation for intensity-calibrated intervention strategies^40,41^.

### Spatially resolved perturbations uncover region-specific regulation in cardiac development

To evaluate the capability of DynPerturb in spatial transcriptomic contexts, we applied DynPerturb to a developmental cardiac datasets to elucidate regionally specific regulatory roles of developmental signals during cardiac morphogenesis. This dataset contains four key stages of entire heart from late fetal to early postnatal development (E20, P01, P04, P14), with anatomical regions annotation (left ventricle, right ventricle, atria) and cell identities annotation (ventricular cardiomyocytes (vCM), atrial cardiomyocytes (aCM), and fibroblasts (FB)) (**Fig. 6A; Supplementary Fig. 17A**,17D). Relative to mature atrial cardiomyocytes, ventricular cardiomyocytes display elevated proliferation and undergo marked expansion and remodeling^44,45^, creating region-specific growth-factor responsiveness. We therefore examined the complementary developmental regulators *Igf2*, *Plagl1* and *Wt1*. *Igf2* is a a key paracrine driver of ventricular proliferation^46^, highly expressed in vCMs from E20 to P01 and rapidly downregulated after P04 (**Fig. 6B**). *Plagl1* in epicardial fibroblasts activates *Igf2*, whereas *Wt1* sustains epicardial signaling^46^. Their concerted, transient embryonic program yields to postnatal homeostasis (**Fig. 6B; Supplementary Fig. 22A, 26A**).

**Figure 6.**
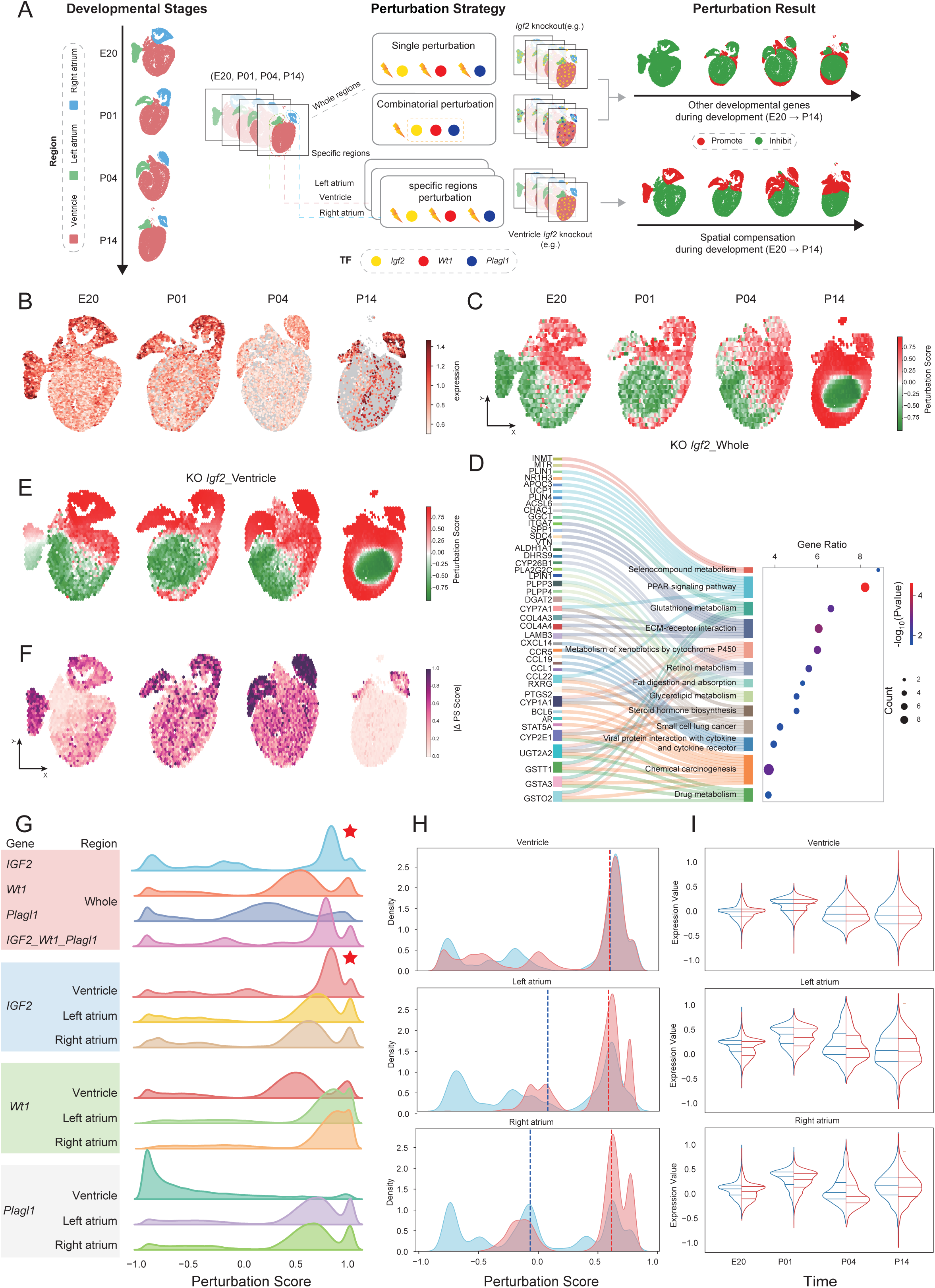
Spatial perturbations demonstrate region-specific regulatory sensitivity during heart development. **(A)** Region-targeted perturbations in the heart, focusing on the left and right atria and the ventricles, with timing varied to assess developmental outcomes. **(B)** Spatial highlight of *Igf2* expression across stages. **(C)** Effects of knocking out *Igf2*, plotted in real spatial coordinates per stage. **(D)** Effects of *Igf2* knockout in the ventricle region specifically. **(E)** Effects of knocking out *Igf2* in ventricle region. **(F)** Absolute difference between full *Igf2* knockout and a combined *Wt1* and *Plagl1* knockout perturbation scores, highlighting regional contributions. **(G)** Ridge plots of overall perturbation scores (PS) for single knockouts of *Igf2*, *Wt1*, or *Plagl1*, combined knockout of all three, and single-gene knockouts restricted to different anatomical regions (ventricle, left atrium, and right atrium). **(H)** Empirical cumulative distribution function (ECDF) of PS scores across regions comparing full *Igf2* knockout (blue) and ventricle-only *Igf2* knockout (red). **(I)** Violin plots comparing node embeddings across regions under full *Igf2* knockout (blue) and ventricle-only *Igf2* knockout (red).

#### Global *Igf2* deletion reveals temporally constrained regulation and developmental cascade effects

We first used DynPerturb to simulate a global knockout of *Igf2*, revealing pronounced temporal and spatial dependencies in its perturbation effects (**Fig. 6C; Supplementary Fig. 18**). *Igf2* is selectively up-regulated in the ventricular myocardium during E20 – P01. Its deletion, simulated by DynPerturb, induces pronounced negative perturbations in the left ventricle and central wall, validating its ventricle-restricted pro-growth function. This implies a diminished developmental potential in the ventricle, while atrial responses only became more intense at P14. This underscores *Igf2*’s critical perinatal role in ventricular development, as its deletion likely suppresses *PI3K/AKT* signaling, reduces mitotic rates, and drives premature cardiomyocyte maturation, ultimately impairing ventricular volume and function, consistent with prior reports^47–50^.

Gene set enrichment analysis (GSEA) further revealed multidimensional pathway disruptions post-*Igf2* loss (**Fig. 6D**): aberrant polyunsaturated fatty acid metabolism and steroid biosynthesis (indicative of premature metabolic maturation, constraining proliferation^48,51^, disrupted ECM-receptor interactions (via *COL4A3*, *LAMB3*, *SDC4*, VTN-compromising cell-matrix adhesion and structural support for ventricular expansion^52^), and *perturbed* retinoic acid metabolism/*cytochrome P450* signaling (impairing signal integration and oxidative stress regulation, exacerbating developmental desynchrony)^53,54^. These convergent disruptions provide a systems-level basis for ventricular defects, most prominent from late gestation to early postnatal life^51^. In contrast, atria showed weaker *Igf2* dependency, with potential postnatal compensation via alternative pathways leading to slower, spatially restricted responses^48,52^.

We further simulated knockouts of *Plagl1* and *Wt1* using DynPerturb. *Plagl1* deletion induced moderate, spatially confined effects, primarily in the subendocardial and subepicardial regions of the ventricle^55^ (**Supplementary Fig. 22-25**). In contrast, *Wt1* knockout produced a broader perturbation pattern, notably affecting the subepicardial mesenchyme and chamber border zones, including the right atrial vicinity^45,50^ (**Supplementary Fig. 26-29**). Simultaneous deletion of *Igf2*, *Plagl1*, and *Wt1* led to a marked amplification of perturbation scores in both central ventricular and subepicardial regions, forming a continuous high-impact domain (**Supplementary Fig. 30**). These results indicate a strong cooperative interaction among these factors in developmental regulation^52,56^ and highlight the essential role of multi-factorial homeostatic control systems in ensuring organ-level robustness during morphogenesis^45,48^.

#### Region restricted knockout uncovers inter chamber signaling and spatial compensation

To further investigate whether localized perturbations could elicit cross regional regulatory feedback, we employed DynPerturb to simulate anatomically precise, region specific *Igf2* knockouts. We focused on ventricle restricted knockout (**Fig. 6E; Supplementary Fig. 19**) as well as left and right atrial restricted deletions (**Supplementary Fig. 20-21**). Compared to global knockout, ventricle restricted knockout produced a substantially weaker negative perturbation scores in the ventricular zone. In addition, the atrial and epicardial compartments exhibited buffering trends, suggesting that neighboring tissues may provide compensatory support via non-cell-autonomous mechanisms to sustain local homeostasis in the face of *Igf2* deficiency-consistent with recently described paradigms of cross regional support mediated by epicardium-mesenchyme-myocardium interactions^45,50,57^. Conversely, in the atrial specific deletions, we also observed significant perturbation responses in the ventricular region. This corroborates prior findings that, although atrial *Igf2* expression is relatively low, it can still influence ventricular development through paracrine signaling^45,57,58^. These regionally constrained perturbation simulations with DynPerturb unveil pronounced spatial crosstalk during early cardiac development: ventricular growth is reliant on supportive cues from the atria, while atrial disturbances may indirectly disrupt ventricular expansion and maturation via feedback mechanisms^45,57^. The spatial inference from DynPerturb supports the central role of anatomical localization in orchestrating cardiac morphogenesis.

#### Whole-heart versus regional perturbations uncover interchamber developmental crosstalk

Then we compared the perturbation responses of global *Igf2* knockout (**Fig. 6C**) with those induced by ventricle restricted deletion (**Fig. 6E**), using DynPerturb simulations. Although both conditions elicited strong perturbation within the ventricular region, global knockout led to more pronounced transcriptional alterations in the atria (**Fig. 6F**), indicating that *Igf2* exerts effects beyond its primary expression domain via inter-chamber signaling. This effect was further corroborated by perturbation score distributions (**Fig. 6G**): while ventricular scores were largely consistent between models, atrial scores under global knockout exhibited a marked leftward shift (**Fig. 6H**), indicative of a broader transcriptomic disturbance. Complementary gene expression analyses revealed dynamic and significant expression changes in atrial regions across multiple time points, in contrast to the relative stability of ventricular profiles (**Fig. 6I**). These findings underscore the non-cell-autonomous support mechanisms between cardiac compartments^44,45^ and highlight the spatial coupling of region specific regulation. *Igf2* appears to mediate developmental coordination through cross regional signaling, aligning with chamber-specific heterogeneity and positional effects reported in recent spatial multi-omic studies of the human heart. Our perturbation-based framework provides a quantitative validation of such cross-compartment regulatory architectures^50,57^.

## Discussion

This study introduces DynPerturb, a time-resolved, multiscale perturbation modeling framework designed to systematically capture the mechanistic effects of gene interventions in dynamic spatiotemporal contexts. By constructing time-resolved dynamic regulatory graphs, DynPerturb explicitly models how perturbations propagate through evolving regulatory networks to drive cell-state transitions and tissue-level remodeling. This approach integrates temporal, spatial, and intensity dimensions, allowing for a nuanced understanding of perturbation effects across biological scales. Importantly, DynPerturb preserves causal relationships, reflecting both the timing of exogenous interventions and the delays inherent in their downstream effects.

Applied to three representative biological systems: human kidney disease, hematopoietic development, and murine cardiac morphogenesis. DynPerturb demonstrates its ability to model regulatory network rewiring, lineage fate transitions, and spatial feedback from molecular to tissue scales. In human kidney disease, it reconstructs proximal-tubule regulatory remodeling and damag-repair trajectories; in hematopoietic development, it resolves stage-specific regulators and branching decisions; and in murine cardiac morphogenesis, it tracks region-specific signaling dynamics and tissue patterning during ventricle-atrium formation. Together, these results indicate that DynPerturb provides a unified framework for spatiotemporally consistent perturbation reasoning across complex developmental and disease systems.

DynPerturb models causal regulation and temporal feedback on real-world time axes. Rather than relying on pseudotime inference, it directly leverages experimentally recorded time to build time-resolved regulatory graphs with higher temporal resolution and stricter causal logic, preserving the causal order of exogenous interventions (e.g., gene perturbations, drug treatments) and the downstream response delays they induce. DynPerturb further incorporates cell annotations as auxiliary supervision during training and jointly optimizes them with the link-prediction objective, strengthening the biological semantic consistency and interpretability of the learned embeddings and enhancing cross-dataset generalization. Finally, DynPerturb flexibly encodes single-gene, gene-set, or combinatorial perturbations at arbitrary time points (T), perturbation intensity (I), and spatial domains (S), enabling multiscale response decomposition from genes to cells to tissues within a unified graph model and providing a rigorous tool and conceptual basis for systems biology and precision intervention strategies.

Overall, DynPerturb exhibits robust modeling capacity, enabling precise learning of regulatory mechanisms from time-series data and inference of perturbation effects across molecular, cellular, and tissue levels. It provides a generative in silico model for mechanism discovery and therapeutic strategy design. In future applications, DynPerturb could be used to build generalized mechanistic atlases from large-scale data while allowing light-weight fine-tuning on individual-specific profiles, enabling precise modeling of personal developmental trajectories and regulatory deviations. This strategy-combining mechanistic priors with individualized adaptation-offers a principled route to accommodate regulatory variation in disease states while preserving systemic architecture. As its capabilities expand, DynPerturb holds promise for developmental dynamics modeling, regulatory mechanism dissection, and rational intervention design-accelerating translation from transcriptional networks to personalized therapeutic strategies.

## Method

### DynPerturb model architecture

DynPerturb is a temporal biological interaction based graph modeling framework. In this framework, each biological relation, including gene regulatory interactions, cell–cell transition probabilities, and spatial adjacency, is encoded as a time-stamped edge and propagated sequentially along the temporal axis to construct an evolving graph structure. Nodes, representing entities such as genes or cells, participate in biological interactions in temporal order and update their internal states upon each interaction, thereby forming a temporally resolved, traceable embedding trajectory that reflects their regulatory history (**Figure 1A;Supplementary Fig. 1A**).

Each node v ∈ V is associated with static features ***x****_v_* ∈ ℝ*^dv^* given by gene expression values.Formally, we construct a dynamic graph *G =*(*V,E,T)*,where each edge is defined as a 5-tuple:

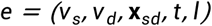

Here, *v_s_* and *v_d_* denote the source and destination nodes, *x_sd_* ∈ ℝ*^de^* represents edge features such as regulatory strength, transition probability, or spatial location, and *t* ∈ ℝ*^+^* denotes the real-valued timestamp derived from the source node. The final element l is a task-specific edge label: for link prediction, *I =1* indicates an observed edge, while *I =0* denotes a randomly sampled negative edge (with 1:1 positive-to-negative ratio);for node classification, *I* encodes the cell type of the source node *v_s_*, but remains attached to the edge rather than as a node attribute. This design enables DynPerturb to flexibly support single-task or multi-task training, while maintaining consistent input structure across objectives. These inputs provide the basis for modeling time-dependent state transitions in the underlying regulatory network.

To learn temporally resolved embedding trajectories, each node is equipped with a recurrent memory vector *m_v_*(*t)*∈ ℝ*^dm^*, representing its internal state at time.When an biological interaction occurs, the source and destination node memories are fused with edge and temporal information to form an interaction message:

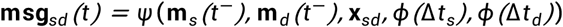

Here, ψ is a learnable message function, and *ф(*· ) encodes elapsed time since the last interaction. The memory is updated via a gated recurrent unit (GRU):

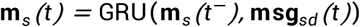

After each update, the model retrieves temporal context from historical neighbors *N_v_(t)={u |(v,u,*∗ *’,t)*∈ *E,t’<t*}, and performs attention-based aggregation:

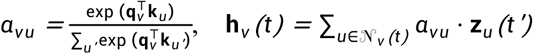

which **q***_v_* and **k***_u_* mean query/key vectors for attention weights, *z_u_(t)*means embedding of neighbor node u at historical time *t*’, *a_vu_* means attention weight assigned to node u by node v and **h***_v_(t)*means updated embedding of node v at time t.

This mechanism allows the model to prioritize temporally and structurally relevant interactions and form causally consistent representations over time.

During training, the model jointly optimizes two objectives to learn the temporal evolution of node states: (1) a link prediction task to capture the dynamic formation of regulatory or intercellular connections, and (2) a node classification task to guide the embedding space toward biologically meaningful structure. The link prediction task is unsupervised and relies on observed interactions and negative sampling to construct training pairs, formulated as:

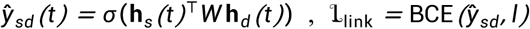

which **h***_s_(t)*and **h***_d_(t)*means embeddings of source and destination nodes at time t and **W***_ℎ_* means projection matrix.

The node classification task incorporates biological annotations (e.g., cell type or function) as supervisory labels and is defined as:

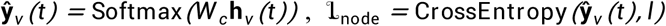

which **W***_c_* means classification weight matrix and **h***_v_(t)*means embedding of node v.

The overall loss combines both objectives with a weighting factor h :

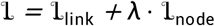

Training is conducted strictly in chronological order of the interactions. The core loop is outlined as follows:

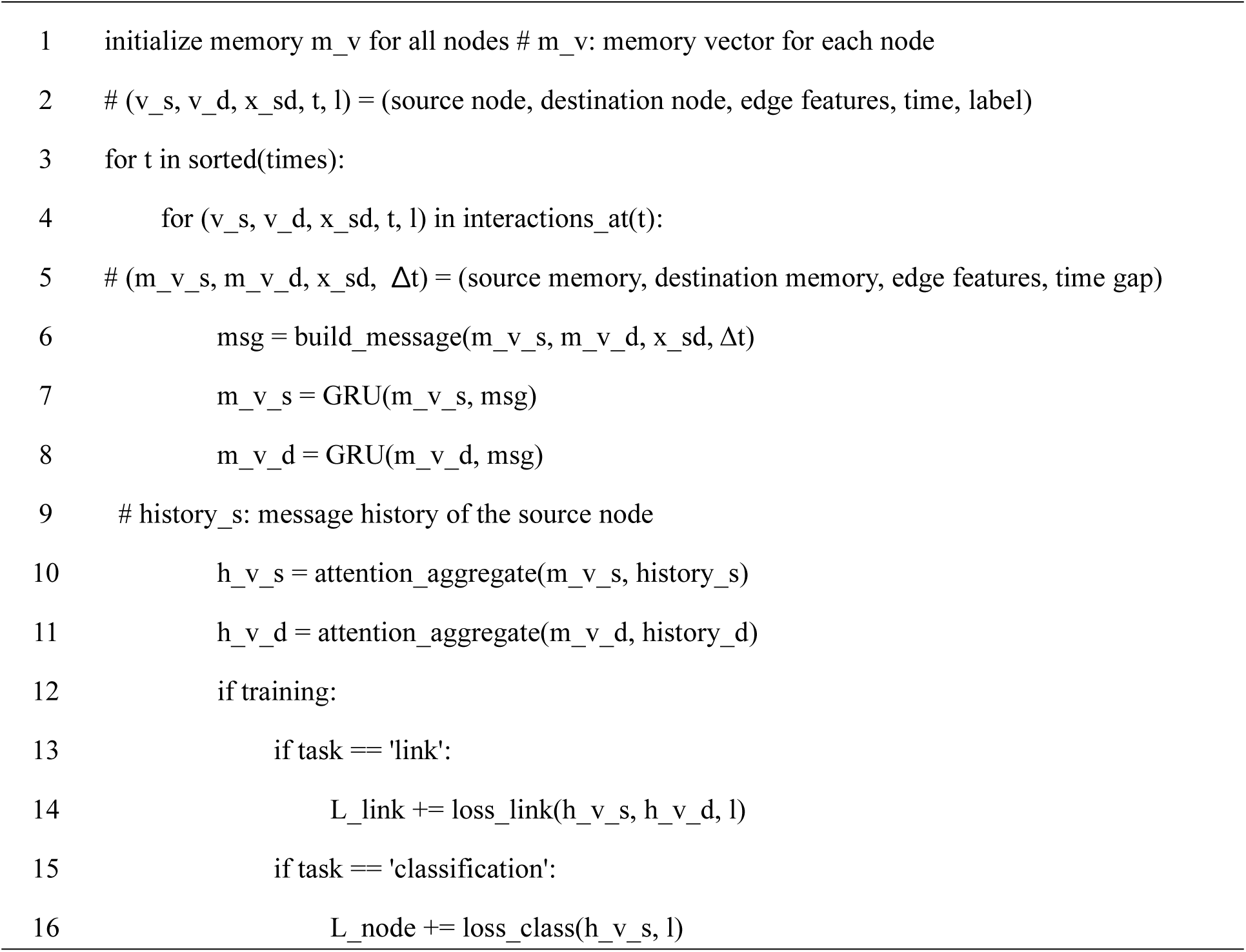

After training, we perform forward inference to extract temporally resolved node embeddings. Whether the input consists of the original dataset or perturbed data, DynPerturb operates on the same sequence format. The model sequentially updates node memories and aggregates neighborhood context at each interaction timestamp to compute node representations. This inference procedure is purely feedforward and does not involve gradient computation or weight updates.

The core embedding extraction process is outlined below:

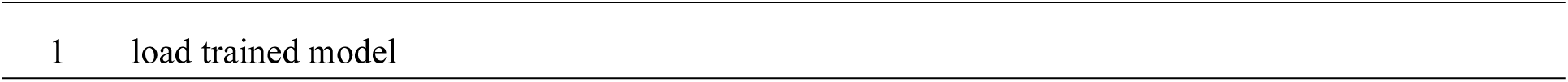
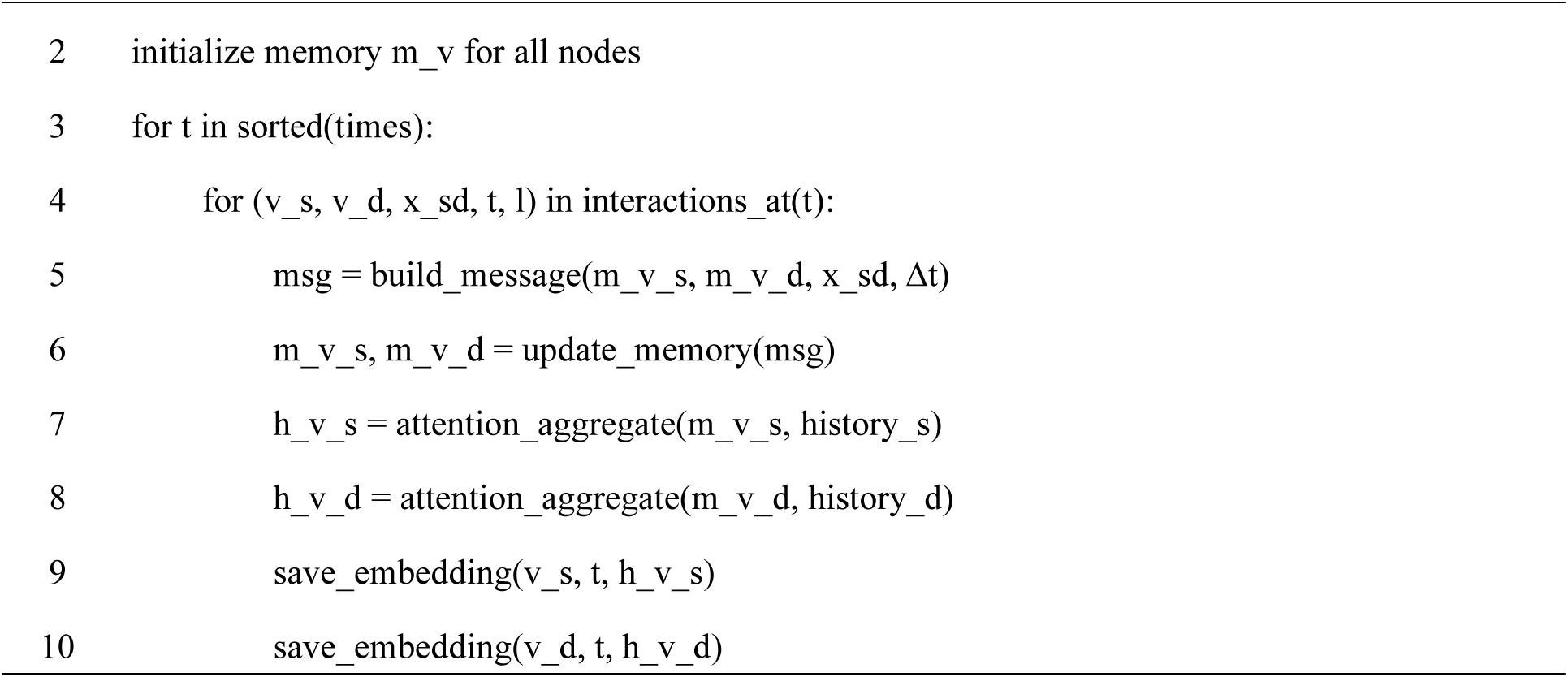

These embeddings are subsequently used for a range of downstream analyses involving perturbations at multiple biological scales, including gene-level regulatory disruptions, cell state reprogramming, and spatially resolved perturbation effects (**Figure 1B**).

### Methodological advances

DynPerturb was developed to address key challenges in modeling gene perturbations within dynamic biological systems. It introduces two major methodological innovations designed to more faithfully capture complex regulatory behaviors over time and across spatial contexts.

First, DynPerturb explicitly models regulatory processes along true biological time by employing a dynamic graph framework driven by biological interactions. Unlike static graph-based methods, it does not rely on a fixed, pre-defined network topology and can accommodate structural rewiring as cell states evolve. In contrast to pseudotemporal approaches, DynPerturb operates directly on time-stamped biological interaction sequences rather than inferring trajectories from similarity-based ordering. This design preserves temporal context and causal ordering of perturbations, enabling the model to capture biologically realistic phenomena such as feedback regulation, cascading amplification, and discrete state transitions.

Second, DynPerturb integrates cell-type annotations as supervisory signals to guide the learning of biologically meaningful embeddings. Built on unsupervised structural modeling, this lightweight semantic constraint aligns node representations with known cellular identities through an auxiliary classification task. This improves the model’s ability to resolve phenotypic boundaries, which is particularly important in heterogeneous systems such as developmental lineages or pathological transitions. Although it relies on minimal prior knowledge, with cell type as the only external input, this supervision significantly enhances the biological coherence of the learned embedding space and improves performance in downstream analyses.

Third, DynPerturb flexibly supports a wide spectrum of perturbation schemes, including different perturbation types (knockout, overexpression, downregulation), varied intervention timings along the biological trajectory, graded perturbation strengths, sustained versus transient perturbations, and combinatorial gene targeting. This design enables systematic exploration of how timing, dosage, duration, and multi-gene interactions shape regulatory dynamics, allowing the model to capture realistic intervention scenarios and predict complex perturbation outcomes.

Together, these innovations enable DynPerturb to overcome the limitations of static and pseudotemporal models by combining structural and semantic learning in a synergistic manner. Even with limited prior information, it generates robust, interpretable embeddings that are well-suited for modeling perturbations across multiple biological scales and contexts.

### Training parameters

During training, DynPerturb processes input as a temporally ordered stream of biological events, ensuring that node embeddings reflect biologically realistic causal progression. Events are strictly partitioned in chronological order: 70% for training, 15% for validation, and 15% for testing, preserving temporal causality throughout.

Training is performed using the Adam optimizer (learning rate of 1×10⁻⁴), with a dropout rate of 0.1 and batch sizes of 64, depending on dataset size. All modules employ 100-dimensional fused structural temporal encodings, and the final node embeddings are configurable in dimension, with a default size of 1,000. Temporal neighbors are aggregated using a single-layer, dual-head attention mechanism. For link prediction, negative edges are sampled at a 1:1 ratio to positive edges. When node classification is included, it is trained jointly with link prediction under an equal loss weighting scheme (1:1).

DynPerturb optionally activates a global memory module to capture long-range temporal dependencies. Training employs early stopping based on validation performance, and inference is performed strictly chronologically with frozen weights. The final model selected for downstream tasks corresponds to the epoch with the best validation metrics.

Details regarding the CPUs and GPUs used for model training are provided in **Supplementary Fig. 2**.

### Datasets and Preprocessing

Four time-series transcriptomic datasets were downloaded from https://zenodo.org/records/6720690#.YrXQjHZBz4Y, including two hESC systems and two mESC systems. A chronic kidney disease single-cell transcriptomic dataset was obtained from cellxgene, containing 7,853 proximal tubule cells, 24,113 genes, and 15 time points across individuals aged 30 to 66 years. A hematopoietic single-cell transcriptomic dataset was also obtained from cellxgene, comprising 88,861 cells, 28,121 genes, and 26 time points covering donor ages from 19 to 84 years, with cell types consolidated into eight major categories. In addition, a mouse cardiac development spatial transcriptomic dataset was downloaded from https://www.ncbi.nlm.nih.gov/geo/query/acc.cgi?acc=GSE178636, consisting of 48,909 spatial bins, 23,397 genes, and four developmental stages (E20, P01, P04, P14).

For all datasets, preprocessing focused on selecting the relevant cell types of interest, followed by batch correction across different time points to minimize technical variation. Expression matrices were normalized and highly variable genes were selected, after which dimensionality reduction was performed using principal component analysis. All procedures were conducted using the default parameters implemented in Scanpy and Harmony.

### Graph contruction

DynPerturb provides a generalizable framework to simulate time-varying biological processes, learn temporal dynamics, and build robust models for subsequent perturbation inference. DynPerturb requires input graphs that encode both temporal dynamics and biologically meaningful structure. Each dynamic graph defines nodes representing genes, cell states, or other biological entities, and edges capturing regulatory, developmental, spatial, or other context-specific relationships that evolve over time. These graphs serve as structured representations of context-specific regulatory programs across conditions or stages. We constructed four representative types of dynamic graphs, each tailored to the characteristics of a specific dataset:

For benchmarking analyses, graphs were generated using the CellOracle^3^ framework, which infers gene regulatory networks from single-cell data. A genome-wide base regulatory network was first assembled from chromatin accessibility information and transcription factor motif annotations, and subsequently refined with expression data across time-resolved subsets. Only regulatory edges that were supported as true interactions were retained during filtering. Each interaction was encoded with information on the transcription factor, its target gene, the associated regulatory strength, the relevant time point, and a binary indicator reflecting the validity of the edge. Genes served as the nodes of the benchmarking graphs, and their feature vectors were derived from sctransform-normalized expression profiles.

For the chronic kidney disease dataset, dynamic gene regulatory networks were reconstructed using CellOracle^3^. A genome-wide based network was generated from chromatin accessibility and transcription factor motif information, and subsequently re-weighted using expression data for each of the 15 time points. Statistically significant edges were retained to produce time-resolved GRNs, which were then subdivided by proximal tubule cell clusters. Each regulatory interaction was encoded as a quintuple including the source transcription factor, target gene, interaction strength, time point, and binary label. Genes served as graph nodes, with feature vectors derived from normalized expression profiles of 1,000 sampled cells per cluster per time point.

For the hematopoietic dataset, graphs were constructed to represent probabilistic differentiation trajectories using Palantir^59^, which models cell fate probabilities and pseudotime from single-cell transcriptomic data. Multipotent progenitor cells from the earliest donor were defined as the root state, and Palantir was applied to infer pseudotime ordering and transition probabilities between cells. Directed edges with transition probabilities greater than 0.1 were retained, capturing biologically meaningful developmental transitions while maintaining graph sparsity. Each edge was encoded as a five-element tuple containing the source cell, target cell, transition probability, timestamp, and source type label. Graph nodes represented individual cells, with features defined by log-normalized expression of the top highly variable genes.

For the cardiac spatial transcriptomic dataset, dynamic spatial graphs were constructed to capture both tissue adjacency and temporal progression. Each spatial bin was treated as a node, and edges were defined using an eight-neighborhood adjacency structure in which bins were connected to directly adjacent neighbors along both orthogonal and diagonal directions. Each edge was encoded as a quadruple containing the source bin, target bin, developmental stage, and type label. Developmental time points were discretized into numerical values, and node features were defined by expression profiles of highly variable genes. This representation preserved spatial topology, gene expression, and temporal information within the same graph structure.

### Benchmarking

We systematically evaluated DynPerturb across multiple public datasets to determine its ability to model dynamic regulatory systems. The evaluation was designed to test whether the model could faithfully capture the temporal evolution of node states and generate structurally consistent, biologically meaningful embeddings for downstream analysis. For benchmarking, each dataset was randomly split into training, validation, and test sets at a ratio of 70:15:15. Gene regulatory network (GRN) prediction was conducted on the training sets, and performance was assessed on the held-out test sets. Prediction accuracy was quantified using the area under the receiver operating characteristic curve (AUC), which was calculated by plotting the true positive rate (TPR) against the false positive rate (FPR) across thresholds.

In the link prediction task, DynPerturb was tested on four widely used GRN benchmarks, including two human and two mouse datasets. This task evaluates whether the model can infer future regulatory interactions from historical graph structures. DynPerturb consistently outperformed baseline methods, including TDL (Temporal Deep Learning), dynDeepDRIM (Dynamic Deep Recurrent Interaction Model), and MI (Mutual Information), as well as PCC (Pearson Correlation Coefficient), achieving higher AUC scores and related metrics. TDL and dynDeepDRIM incorporate dynamic modeling approaches to capture temporal dependencies, while MI and PCC are static correlation-based measures that do not account for temporal variation in regulatory interactions. These results highlight DynPerturb’s ability to capture both the formation and temporal propagation of regulatory interactions under complex time-dependent conditions (Figure 2A-B).

We then examined the biological interpretability and semantic coherence of the learned embeddings through a node classification task (Supplementary Figure 7A). In human bone marrow development data, DynPerturb accurately predicted cell identities across multiple hematopoietic lineages. Most misclassifications occurred between phenotypically similar or developmentally adjacent populations, such as LMPP, MPP, and Cycling Progenitors. These “semantically adjacent” errors suggest that the learned embeddings preserve biological relationships and reflect developmental continuity in latent space.

To further evaluate spatial generalizability, we applied DynPerturb to a spatial transcriptomics dataset of mouse heart tissue. The model successfully resolved functionally distinct subpopulations, including aCM, vCM, and FB, despite considerable spatial heterogeneity (Supplementary Figure 17A). The normalized confusion matrix confirmed strong classification accuracy and spatial resolution, demonstrating that DynPerturb retains semantic separability while effectively integrating anatomical context.

Overall, DynPerturb shows robust performance across diverse data modalities. It not only captures the dynamic assembly of regulatory interactions but also learns biologically meaningful embeddings, providing a strong foundation for downstream applications such as perturbation simulation and trajectory analysis.

### Perturb Analysis

#### Embedding Difference Vector

For each perturbation scenario, node embeddings were first computed under both unperturbed and perturbed conditions. The perturbation effect for each gene or cell was represented as the embedding difference vector Δℎ *=*ℎ^pert^ − ℎ^base^, where ℎ^pert^ and ℎ^base^ denote the embeddings in the perturbed and baseline conditions, respectively. The magnitude of this difference was used to quantify the strength of perturbation-induced changes.

#### Mean Expression Difference

The Mean Expression Difference refers to the average of the Embedding Difference Vectors across all samples.

#### Perturbation Score (PS)

CellOracle^3^ proposed a strategy to assess the directionality of perturbation effects by comparing embedding shifts to the baseline developmental vector field. For each node i, the score was defined as:

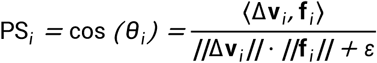

where Δv_i_ is the perturbation-induced embedding shift, **f***_i_* is the baseline developmental direction vector, and c is a small constant to avoid division by zero. Values close to 1 indicate reinforcement of the baseline trajectory, values near – 1 indicate reversal, and values around 0 indicate ambiguous shifts.

### Sum of Negative Perturbation Scores

Sum of Negative Perturbation Scores was calculated by summing all perturbation scores (PS) less than zero, followed by normalization. The resulting values were visualized using a scatter plot, where proximity to a given axis indicates stronger suppression of the corresponding lineage. This analysis was adapted from the CellOracle^3^ framework.

### ECDF analysis

Empirical cumulative distribution functions (ECDF) were used to evaluate the global distribution of perturbation metrics across cell populations. For each metric, the empirical distribution function was calculated as:

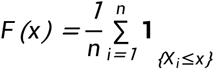

where *X_i_* are the observed values across nnn nodes or cells. ECDF curves allowed direct comparison of perturbation responses between clusters, time points, or experimental conditions, highlighting heterogeneity and population-level shifts.

Δ cosine. To further quantify perturbation outcomes at the regulatory network level, we first computed the temporal mean embedding for each gene across all time points, resulting in a gene-by-time representation matrix. This matrix captures the dynamic behavior of each gene throughout the time course. To extract higher-order structural information, we input this matrix into a two-layer Graph Convolutional Network (GCN) trained on the corresponding cluster-specific temporal GRN.The GCN learned compressed graph-level embeddings that encapsulate the overall regulatory state of the system under both perturbed and baseline conditions. We then computed Δ cosine, defined as the cosine distance between perturbed and control graph embeddings, to quantify network-level differences.

## Data availability

The four embryonic stem cell (ESC) datasets for benchmarking were obtained from reference^14^. These datasets can be accessed at https://zenodo.org/records/6720690#.YrXQjHZBz4Y. The human kidney dataset was obtained from reference^20^. This dataset can be accessed at https://cellxgene.cziscience.com/e/dea717d4-7bc0-4e46-950f-fd7e1cc8df7d.cxg/. The human hematopoietic dataset was obtained from reference^60^. This dataset can be accessed at https://cellxgene.cziscience.com/e/cd2f23c1-aef1-48ae-8eb4-0bcf124e567d.cxg/. The mouse heart spatial transcriptomics dataset was obtained from reference^61^. This dataset can be accessed at https://www.ncbi.nlm.nih.gov/geo/query/acc.cgi?acc=GSE178636 (**Supplementary Table 2**). The training data and model parameters used in this study are available at Figshare (DOI: 10.6084/m9.figshare.30040642).

## Code availability

The DynPerturb model and all related analysis code can be accessed athttps://github.com/BGIResearch/DynPerturb.

## Acknowledgement

We acknowledge the Stomics Cloud platform (https://cloud.stomics.tech/) for providing GPU computational resources. We thank the colleagues in our research group for inspiring discussion and their contributions.

## Author contributions

S.F., Y.Z., and Y.L. conceptualized the study. H.Q. and Y.Z. were responsible for model design and tool implementation. H.Q., Y.Z., Z.G., and L.H. performed data analysis and model evaluation. S.F., Y.L., Q.L., C.L., T.X., and Z.D. contributed key ideas and advice. H.Q. wrote the manuscript. S.F., Y.Z., and Y.L. supervised the study.

---

**Supplementary Figure 1. Ablation analysis of model performance in the mESC1 dataset.**

(A) Model Architecture. The model constructs dynamic graphs from time-series features and uses a temporal GNN to learn node embeddings for tasks like link prediction and node classification.

(B) Ablation analysis with reduced numbers of cells, genes, and time points in the mESC1 dataset.

(C) Ablation study on the number of time points, gene numbers, and highly variable gene (HVG) numbers for link prediction AUC on unseen nodes in the mESC1 dataset.

(D) Scatterplot showing the number of cells and time points in each dataset.

**Supplementary Figure 2. Computational resource usage during DynPerturb execution.**

Resource utilization during a representative simulation run for the three core modules of the model: the Gene Regulatory Network (GRN; blue), Cell Development (red), and Spatial (green) modules.

(A) CPU utilization over time for each module.

(B) GPU utilization over time for each module.

(C) Memory utilization over time for each module.

**Supplementary Figure 3. Cell composition, model evaluation, and regulatory network features in proximal tubule cells.**

(A) Cell type composition over time in proximal tubule cells.

(B) Cluster-wise AUC performance of link prediction.

(C) Training loss and ablation analysis with reduced numbers of cells, genes, and time points.

(D) Top 20 differentially expressed genes in aPT and PT-S1/S2 clusters.

(E) Partial regulatory network of aPT-B and temporal changes in the expression levels of *ELF3*, *KLF6*, and *KLF10*.

**Supplementary Figure 4. Key regulatory genes and perturbation effects in aPT-B.**

(A) Top 100 genes in the aPT-B GRN ranked by degree centrality, in-degree, and out-degree.

(B) Distribution statistics of the four perturbation conditions (aPT KO, PT KO, aPT UP, and PT UP) corresponding to Figure 3C (absolute changes in node embeddings of 40 marker genes: 20 PT markers and 20 aPT markers, separated by a dashed line, after perturbing *ELF3*, *KLF6*, and *KLF10* in both PT and aPT).

(C) Cluster-based calculation of cosine distance changes between PT and aPT, corresponding to Figure 3F. Top:KO in aPT-A and aPT-B (aPT to PT direction); bottom:UP in PT-S1-A/B/C/D and PT-S2-A/B/C (PT to aPT direction).Red indicates a shift closer to the reference state, whereas blue indicates a shift further away.

(D) Band-shaped average line plots corresponding to Figure 4C (node embedding changes after full, quarter (reduced to one-quarter of the original expression level), or half knockout of *ELF3*, *KLF6*, and *KLF10* from the earliest time point.

**Supplementary Figure 5. Combinatorial and temporal effects of ELF3, KLF6, and KLF10 knockouts in aPT-B.**

(A) Embedding dynamics following knockout (KO) of six combinations (individual and pairs) of *ELF3*, *KLF6*, and *KLF10* starting at different time points in aPT-B.

**Supplementary Figure 6. Combinatorial and temporal effects of ELF3, KLF6, and KLF10 perturbations in aPT-A.**

(A) Embedding dynamics following full, quarter (reduced to one-quarter of the original expression level), or half knockout (KO) of *ELF3*, *KLF6*, and *KLF10* from the beginning in aPT-A. The 80 most perturbed genes are shown.

(B) Embedding dynamics following knockout (KO) or upregulation (UP) of *ELF3*, *KLF6*, and *KLF10* starting at different time points in aPT-A. The 80 most perturbed genes are shown.

**Supplementary Figure 7. Evaluation of model performance and ablation analysis in hematopoietic development.**

(A) Training loss curve and ablation results with reduced number of cells, genes, and time points.

**Supplementary Figure 8. Effects of knockout of GM lineage regulatory transcription factors on hematopoietic differentiation.**

(A) Effects of *IRF8* knockout on lineage differentiation.

(B) Effects of *CEBPA* knockout on lineage differentiation.

(C) Effects of *CEBPE* knockout on lineage differentiation.

(D) ECDF plots of PS scores in GM and ME lineages under the conditions in (A–C).

**Supplementary Figure 9. Effects of knockout of ME lineage transcription factors on hematopoietic differentiation.**

(A) Effects of *GATA1* knockout on lineage differentiation.

(B) Effects of *KLF1* knockout on lineage differentiation.

(C) Effects of *GATA2* knockout on lineage differentiation.

(D) Effects of *FLI1* knockout on lineage differentiation.

(E) ECDF plots of PS scores in GM and ME lineages under the conditions in (A–D).

**Supplementary Figure 10. Effects of upregulation of GM lineage transcription factors on hematopoietic differentiation.**

(A) Effects of *IRF8* upregulation on lineage differentiation.

(B) Effects of *CEBPA* upregulation on lineage differentiation.

(C) ECDF plots of PS scores in GM and ME lineages under the conditions in (A, B).

**Supplementary Figure 11. Effects of upregulation of ME lineage transcription factors on hematopoietic differentiation.**

(A) Effects of *GATA1* upregulation on lineage differentiation.

(B) Effects of *KLF1* upregulation on lineage differentiation.

(C) ECDF plots of PS scores in GM and ME lineages under the conditions in (A, B).

**Supplementary Figure 12. Effects of half-strength upregulation of GM lineage transcription factors on hematopoietic differentiation.**

(A) Effects of half-strength upregulation of *IRF8* on lineage differentiation.

(B) Effects of half-strength upregulation of *CEBPA* on lineage differentiation.

(C) Effects of half-strength upregulation of *CEBPE* on lineage differentiation.

(D) ECDF plots of PS scores in GM and ME lineages under the conditions in (A–C).

**Supplementary Figure 13. Effects of half-strength upregulation of ME lineage transcription factors on hematopoietic differentiation.**

(A) Effects of half-strength upregulation of *GATA1* on lineage differentiation.

(B) Effects of half-strength upregulation of *KLF1* on lineage differentiation.

(C) Effects of half-strength upregulation of *GATA2* on lineage differentiation.

(D) Effects of half-strength upregulation of *FLI1* on lineage differentiation.

(E) ECDF plots of PS scores in GM and ME lineages under the conditions in (A–D).

**Supplementary Figure 14. Combined upregulation of A-group and B-group transcription factors on hematopoietic differentiation.**

(A) Effects of simultaneous upregulation of B-group transcription factors (*GATA1, KLF1*) on lineage differentiation.

(B) Effects of simultaneous upregulation of both A-group (*IRF8, CEBPA*) and B-group (*GATA1, KLF1*) transcription factors on lineage differentiation.

(C) ECDF plots of PS scores in GM and ME lineages under the conditions in (A, B).

**Supplementary Figure 15. Effects of half-maximal upregulation of GM lineage transcription factors on hematopoietic differentiation.**

(A) Effects of half-maximal upregulation of *IRF8* on lineage differentiation.

(B) Effects of half-maximal upregulation of *CEBPA* on lineage differentiation.

(C) Effects of half-maximal upregulation of *CEBPE* on lineage differentiation.

(D) ECDF plots of PS scores in GM and ME lineages under the conditions in (A–C).

**Supplementary Figure 16. Effects of half-maximal upregulation of ME lineage transcription factors on hematopoietic differentiation.**

(A) Effects of half-maximal upregulation of *GATA1* on lineage differentiation.

(B) Effects of half-maximal upregulation of *KLF1* on lineage differentiation.

(C) Effects of half-maximal upregulation of *GATA2* on lineage differentiation.

(D) Effects of half-maximal upregulation of *FLI1* on lineage differentiation.

(E) ECDF plots of PS scores in GM and ME lineages under the conditions in (A–D).

**Supplementary Figure 17. Evaluation of model performance and ablation analysis during heart development.**

(A) Spatial (X-Y) plots of cells colored by cell type across four developmental stages (E20, P01, P04, P14).

(B) Training loss and ablation analysis for reduced numbers of cells and genes.

(C) UMAP of cells colored by cell type and developmental stage and Pseudotime trajectory inferred by Palantir.

**Supplementary Figure 18. Igf2 knockout in the whole heart during heart development.**

(A) Spatial perturbation vector fields, PS score, and perturbation intensity under *Igf2* knockout in the whole heart across four stages (E20, P01, P04, P14).

(B) Violin plots of Igf2-related embeddings under *Igf2* knockout in the whole heart.

(C) ECDF plots of PS scores across anatomical regions under *Igf2* knockout.

**Supplementary Figure 19. Igf2 knockout in the ventricle during heart development.**

(A) Spatial perturbation vector fields, PS score, and perturbation intensity under *Igf2* knockout in the ventricle across four stages (E20, P01, P04, P14).

(B) Violin plots of *Igf2*-related embeddings under *Igf2* knockout in the ventricle.

(C) ECDF plots of PS scores in the ventricle under *Igf2* knockout.

**Supplementary Figure 20. Igf2 knockout in the left atrium during heart development.**

(A) Spatial perturbation vector fields, PS score, and perturbation intensity under *Igf2* knockout in the left atrium across four stages (E20, P01, P04, P14).

(B) Violin plots of *Igf2*-related embeddings under *Igf2* knockout in the left atrium.

(C) ECDF plots of PS scores in the left atrium under *Igf2* knockout.

**Supplementary Figure 21. Igf2 knockout in the right atrium during heart development.**

(A) Spatial perturbation vector fields, PS score, and perturbation intensity under *Igf2* knockout in the right atrium across four stages (E20, P01, P04, P14).

(B) Violin plots of *Igf2*-related embeddings under *Igf2* knockout in the right atrium.

(C) ECDF plots of PS scores in the right atrium under *Igf2* knockout.

**Supplementary Figure 22. Plag1 knockout in the whole heart during heart development.**

(A) Spatial highlight of Plag1 expression across stages.

(B) Spatial perturbation vector fields, PS score, and perturbation intensity under Plag1 knockout in the whole heart across four stages (E20, P01, P04, P14).

(C) Violin plots of Plag1-related embeddings under Plag1 knockout in the whole heart.

(D) ECDF plots of PS scores across anatomical regions under Plag1 knockout.

**Supplementary Figure 23. Plagl1 knockout in the ventricle during heart development.**

(A) Spatial perturbation vector fields, PS score, and perturbation intensity under *Plagl1* knockout in the ventricle across four stages (E20, P01, P04, P14).

(B) Violin plots of *Plagl1*-related embeddings under *Plagl1* knockout in the ventricle.

(C) ECDF plots of PS scores in the ventricle under *Plagl1* knockout.

**Supplementary Figure 24. Plagl1 knockout in the left atrium during heart development.**

(A) Spatial perturbation vector fields, PS score, and perturbation intensity under *Plagl1* knockout in the left atrium across four stages (E20, P01, P04, P14).

(B) Violin plots of *Plagl1*-related embeddings under knockout *Plagl1* in the left atrium.

(C) ECDF plots of PS scores in the left atrium under *Plagl1* knockout.

**Supplementary Figure 25. Plagl1 knockout in the right atrium during heart development.**

(A) Spatial perturbation vector fields, PS score, and perturbation intensity under *Plagl1* knockout in the right atrium across four stages (E20, P01, P04, P14).

(B) Violin plots of *Plagl1*-related embeddings under *Plagl1* knockout in the right atrium.

(C) ECDF plots of PS scores in the right atrium under *Plagl1* knockout.

**Supplementary Figure 26. Wt1 knockout in the whole heart during heart development.**

(A) Spatial highlight of *Wt1* expression across stages.

(B) Spatial perturbation vector fields, PS score, and perturbation intensity under *Wt1* knockout in the whole heart across four stages (E20, P01, P04, P14).

(C) Violin plots of *Wt1*-related embeddings under *Wt1* knockout in the whole heart.

(D) ECDF plots of PS scores across anatomical regions under *Wt1* knockout.

**Supplementary Figure 27. Wt1 knockout in the ventricle during heart development.**

(A) Spatial perturbation vector fields, PS score, and perturbation intensity under *Wt1* knockout in the ventricle across four stages (E20, P01, P04, P14).

(B) Violin plots of *Wt1*-related embeddings under *Wt1* knockout in the ventricle.

(C) ECDF plots of PS scores in the ventricle under *Wt1* knockout.

**Supplementary Figure 28. Wt1 knockout in the left atrium during heart development.**

(A) Spatial perturbation vector fields, PS score, and perturbation intensity under *Wt1* knockout in the left atrium across four stages (E20, P01, P04, P14).

(B) Violin plots of *Wt1*-related embeddings under *Wt1* knockout in the left atrium.

(C) ECDF plots of PS scores in the left atrium under *Wt1* knockout.

**Supplementary Figure 29. Wt1 knockout in the right atrium during heart development.**

(A) Spatial perturbation vector fields, PS score, and perturbation intensity under *Wt1* knockout in the right atrium across four stages (E20, P01, P04, P14).

(B) Violin plots of *Wt1*-related embeddings under *Wt1* knockout in the right atrium.

(C) ECDF plots of PS scores in the right atrium under *Wt1* knockout.

**Supplementary Figure 30. Combined knockouts of Igf2, Plag1, and Wt1 during heart development.**

(A) Spatial perturbation vector fields, PS score, and perturbation intensity under combined knockout of *Igf2*, *Plag1*, and *Wt1* across four stages (E20, P01, P04, P14).

(B) Absolute PS score differences between full *Igf2* knockout and combined knockout.

(C) Violin plots of *Igf2*-, *Plag1*-, and *Wt1*-related embeddings under combined knockout.

(D) ECDF plots of PS scores across different anatomical regions under combined knockout of *Igf2*, *Plag1*, and *Wt1*.

